# The role of Med15 sequence features in transcription factor interactions

**DOI:** 10.1101/2024.05.04.592524

**Authors:** David G. Cooper, Shulin Liu, Emma Grunkemeyer, Jan S. Fassler

## Abstract

Med15 is a general transcriptional regulator and subunit within the tail module of the RNA Pol II Mediator complex. The *S. cerevisiae* Med15 protein has a well-structured N-terminal KIX domain, three Activator Binding Domains (ABDs), several naturally variable polyglutamine (poly-Q) tracts (Q1, Q2, Q3) embedded in an intrinsically disordered central region, and a C-terminal Mediator Association Domain (MAD). We investigated how the presence of ABDs and changes in length and composition of poly-Q tracts influences Med15 activity and function using phenotypic, gene expression, transcription factor interaction and phase separation assays of truncation, deletion, and synthetic alleles. We found that individual Med15 activities were influenced by the number of activator binding domains (ABDs) and adjacent polyglutamine tract composition. Robust Med15 activity required at least the Q1 tract and the length of that tract modulated activity in a context-dependent manner. We found that loss of Msn2-dependent transcriptional activation due to Med15 Q1 tract variation correlated well with a reduction in Msn2:Med15 interaction strength, but that interaction strength did not always mirror the propensity for phase separation. We also observed that distant glutamine tracts and Med15 phosphorylation affected the activities of the KIX domain, suggesting that intramolecular interactions may affect some Med15-transcription factor interactions. Further, two-hybrid based interaction studies revealed intramolecular interactions between the N-terminal KIX domain and the Q1R domain of Med15.

**Author Summary:** Glutamine tracts are relatively uncommon, but are a feature of many transcriptional regulators including the Med15 subunit of the Mediator Complex which is a large protein complex that plays an important role in gene expression in eukaryotic organisms including yeast and animals. Strains lacking Med15 are compromised in their ability to grow on many kinds of media, under stress conditions, and in fermentation, reflecting its importance in gene expression. Naturally occurring yeast strains specialized for growth in specific environments (e.g., wine, beer, clinical) vary in their glutamine tract lengths, suggesting that the length of glutamine tracts may influence Med15 function in a manner that is adaptive for a specific environment. In this study, we intentionally manipulated the length of the glutamine tracts in Med15 and found that these changes have subtle effects on Med15 interactions with transcription factors, target gene expression and growth. Taken together, our data suggests that glutamine tracts do not themselves mediate critical interactions with partner proteins, but instead may influence the shape of the Med15 protein, thus indirectly affecting the nature of these interactions.

## Introduction

Polyglutamine (poly-Q) tracts, or consecutive repeats of the amino acid glutamine, are a dynamic protein feature with the potential to cause disease by expansion of the underlying microsatellite repeats [1-3]. However, poly-Q tracts have been ascribed non-pathogenic roles that include providing flexible intradomain spacing and protein-protein interaction surfaces [4-8]. Perhaps not surprisingly many of the proteins containing poly-Q tracts are transcriptional regulators [9]. Polyglutamine tract length differences encoded by naturally occurring alleles of genes have been shown to modulate protein activity. Variation in the length of the poly-Q tract of the Clock protein corresponds to alterations in barn swallow breeding [10]. The subcellular localization of Angustifolia in *Populus tremula* is altered by poly-Q tract length changes [11]. Variation in the length of the poly-Q tract of the Notch protein alters developmental phenotypes in fruit flies [12]. Natural and synthetic Q and QA repeat variability in yeast Cyc8 has broad transcriptomic effects [13]. We hypothesize that similarly variable poly-Q tracts in yeast Med15 alter the transcriptional activation function of the protein.

Poly-Q tracts could influence activity by affecting protein-protein interactions, either directly or indirectly; providing disorder to allow a larger set of transient interactions; providing necessary spacing between functional domains; or by providing flexibility to the protein. Poly-Q proteins have a higher number of interactions than other proteins and poly-Q proteins are often found in protein complexes [9]. One interaction surface mediated by poly-Q tracts are coiled-coil domains [14]. In yeast, homodimerization of the Nab3 component of the Nrd1-Nab3 transcription termination complex is facilitated by the coiled-coil structure of a C-terminal Q-tract [5, 15]. The polyglutamine content in the human Med15 ortholog has been shown to form a coiled-coil structure [16]. However, how the multiple poly-Q tracts in yeast Med15 may influence transcriptional activation has not been thoroughly explored.

As a subunit of the RNA Polymerase II Mediator complex, Med15 functions as an interaction hub with other transcriptional regulators to regulate gene expression. Distributed throughout the Med15 sequence are residues and domains that have been shown to correspond to the interactions between Med15 and other proteins. The C-terminal Mediator Association Domain (MAD (aa 799-1081)) permits Med15 to assemble into the Mediator complex [17, 18]. Specifically, conserved amino acid residues 866-910 in MAD are required for Med15 incorporation into Mediator [19, 20]. The domains or regions of Med15 required for interactions with yeast transcription factors (TFs) including Oaf1, Pdr1/3, Msn2, Gcn4, and Gal4 have been mapped with varying resolution [21]. The N-terminal KIX domain (aa 6-90) mediates interactions between Med15 and the Pdr1/3 and Oaf1 TFs [22, 23]. The interactions with other TFs require sequences within the glutamine-rich central region of Med15 and are discussed below.

Med15, formerly Gal11, was initially discovered in conjunction with its role in galactose metabolism [19, 24]. Med15 regulates the expression of galactose metabolism genes through interactions with Gal4 bound to UAS_G_ motifs upstream of *GAL* genes [25]. Another well-studied Med15 interactor is Gcn4. Gcn4 regulates many genes in yeast, including those involved in amino acid biosynthesis [26, 27]. The Gcn4 TF interacts with residues in 4 regions of Med15, the N-terminal KIX domain and three so-called Activator Binding Domains (ABD1 (aa 158-238), ABD2 (aa 277-368), and ABD3 (aa 496-630)) embedded in the intrinsically disordered and glutamine-rich midsection of the protein [20]. Gcn4 and Gal4 interactions with Med15 are described as “fuzzy” meaning that the interactions can be mediated using different subsets of the available hydrophobic residues in interaction pockets [28, 29]. Interacting Gcn4 and Med15 proteins form liquid droplets [30]. Liquid droplet phase separation may be one way to increase the concentration of interactors to compensate for “transient” interactions.

Med15 also interacts with the core environmental stress response transcription factors Msn2/4 to regulate gene expression of stress response genes [31]. Med15 regulates the expression of Msn2 dependent genes *HSP12* and *TFS1* [31]. Although Med15 is not known to interact directly with Hsf1, expression of some heat shock proteins including *SSA4* and *HSP104* is also regulated by *MED15*.

The central section of Med15 (∼aa 100-700) is enriched in amidic amino acids glutamine (16% of residues) and asparagine (11% of residues) [21]. Within this region there are three variable polyglutamine (poly-Q) tracts (longer than 10 residues) which we refer to as Q1 (aa 147-158), Q2 (aa 417-480), and Q3 (aa 674-696) that flank, but are not included in the ABD regions. Q1 and Q3 are simple poly-Q tracts while Q2 consists of a repeated glutamine-alanine motif. While shorter stretches of glutamine are present in Med15, Q1, Q2 and Q3 stand out because they vary in length (number of consecutive Q or QA residues) in *MED15* alleles from different strains of *S. cerevisiae* [21]. Recent studies have addressed the functional implications of naturally occurring tract length variant alleles of *MED15* on resistance to the coal-cleaning chemical, 4-methylcyclohexane methanol [32], and grape juice fermentation [33]. Here we investigate which additional activities of Med15 require these glutamine-rich regions, and the extent to which variations to the polyglutamine tracts in Med15 influence these protein activities.

## Results

The ability of yeast to respond to stress and changes in growth conditions requires the expression of target genes dependent on interactions between Med15 and gene-specific transcription factors. To test the extent to which distinct transcription factor interacting regions and the intervening sequences in Med15 contribute to its activity, we analyzed a series of internal deletions and truncated *MED15* alleles (Fig. 1A) for phenotypes exhibited by the deletion mutant (Fig. 1B). All *MED15* alleles were driven by the native *MED15* promoter and exhibited levels of expression comparable to or greater than wild type (Fig. S1).

**Figure 1.**
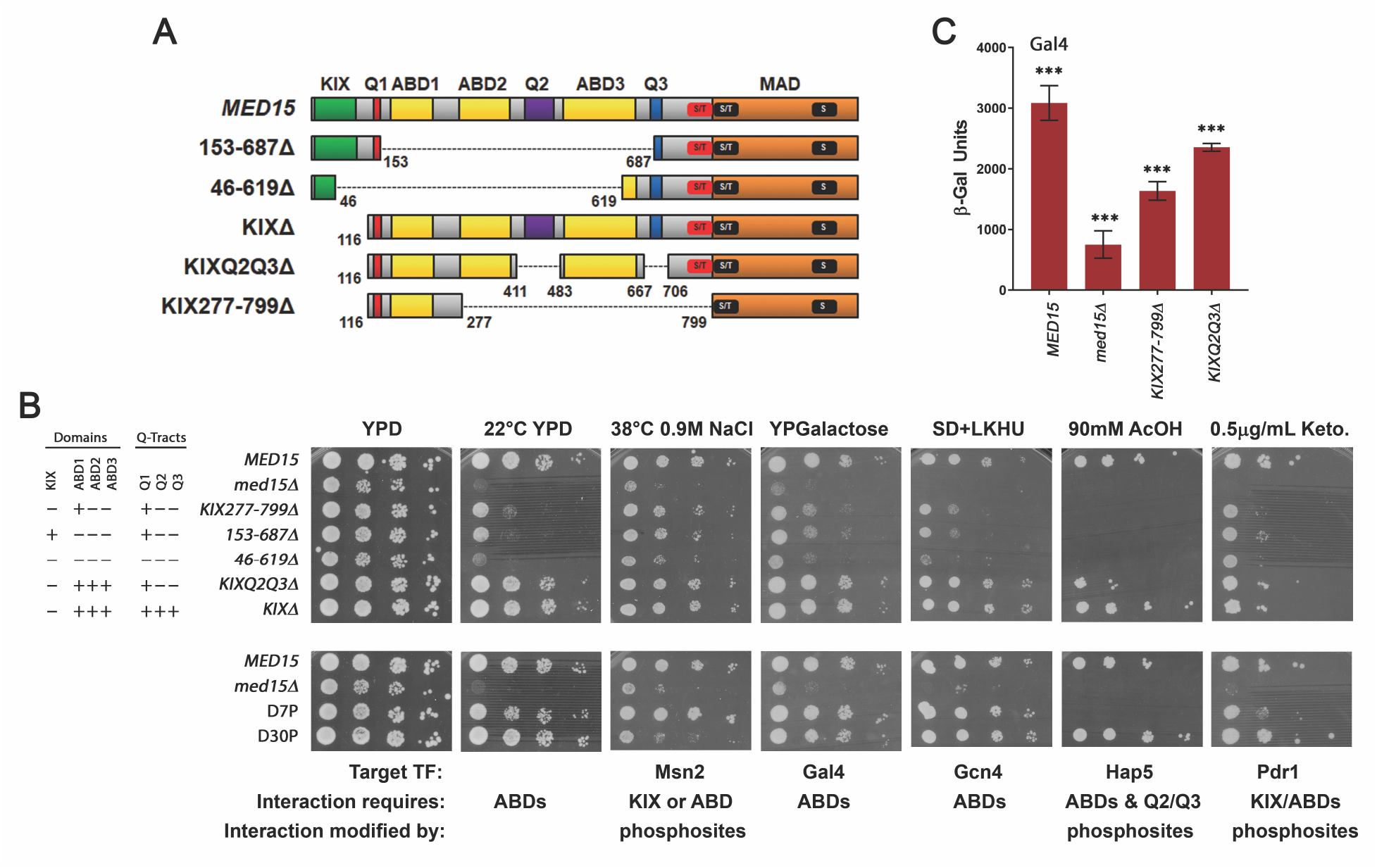
Internal deletions and associated Med15 activities. **(A)** Full length Med15 (1081 aa) domains: KIX (Kinase-inducible domain Interacting Domain), ABD1-3 (Activator Binding Domains), Q1-3 (polyglutamine tracts), MAD (Mediator Association Domain). Regions containing phosphorylation sites (S/T and S) including those dynamically phosphorylated in response to salt stress (red box) and others (black boxes). Numbers represent amino acid residues. KIXΔ-*MED15* constructs lack the KIX domain. KIXQ2Q3Δ-*MED15* constructs lack the KIX domain and the poly-Q tracts for Q2 and Q3. In addition to the construct with the lab allele 12 glutamine residues at Q1, KIXQ2Q3Δ-*MED15* constructs include a 0-Q1 construct as well as versions with different Q1 tract lengths (#-Q1) and non-glutamine sequences. KIX277-799Δ-*MED15* constructs lack the KIX domain and central Q2/Q3 containing region. **(B)** Log-phase cultures of the *med15Δ* strain (OY320) carrying a CEN plasmid with the indicated *MED15* allele, or strains with integrated phospho-site mutations (D7P and D30P) were serially diluted and spotted on YPD media with or without supplements and incubated at 30°C or alternative temperature as specified (38°C or room temperature (22°C)). **(C)** β-galactosidase activity of a Gal4-dependent (UAS_G_-*lacZ*) reporter in a *med15Δ gal80Δ* strain (JF2631) carrying a CEN plasmid with the indicated *MED15* allele. Data points are the averages of at least 3 biological replicates (transformants) and error bars are the standard deviation of the mean. Significant differences were determined using ANOVA analysis with a Tukey post-hoc test, *** p≤0.001.

### Four Med15 Activator Binding Domains contribute additively to Med15 activity

Two internal deletion constructs (153-687Δ and 46-619Δ) lacking most of the central region of Med15 displayed different levels of activity when expressed in a *med15Δ* strain (Fig. 1). Deletion 46-619Δ disrupts or lacks all characterized activator binding domains (ABD or KIX), while deletion 153-687Δ still maintains an intact KIX domain (Fig. 1). Both constructs contain the Mediator Association Domain (MAD), which is necessary for Med15 activity [17, 19]. The 46-619Δ construct failed to fully complement most of the tested *med15Δ* defects (Fig. 1B). Inclusion of the KIX domain, as in the 153-687Δ construct, was sufficient for partial to full complementation of the tested phenotypes except for acetic acid sensitivity (Fig. 1B).

### The ABD1 Region Alone (Q1R) Partially Complements *med15Δ* Defects

KIX277-799Δ-*MED15* retains the glutamine-rich region of Med15 around Q1 (Q1R; aa 116-277) and the MAD domain (aa 799-1081) (Fig. 1A) but lacks the KIX domain, ABD2, ABD3, Q2 and Q3. When expressed in a *med15Δ* strain, KIX277-799Δ-*MED15* partially complemented many of the *med15Δ* phenotypes tested, including utilization of galactose as a carbon source and tolerance to ethanol (not shown) (Fig. 1B), and provided substantial complementation of growth in the presence of combined osmotic and thermal stress.

Only one of the two regions of the Med15 protein involved in interactions with the Gal4 transcription factor (aa 116-176 and aa 566-618) [34] is included in the KIX277-799Δ-*MED15* construct. The presence of this one region (aa 116-176) allowed for partial activation of a Gal4-dependent reporter (Fig. 1C). The reduced level of expression relative to full length *MED15* is consistent with the phenotype of KIX277-799Δ-*MED15* on galactose plates (Fig. 1B).

### KIX domain activity

Specific activities have been attributed to the discretely-folded and well-conserved KIX domain (aa 6-90), including interactions with the transcription factor Oaf1 to regulate the expression of fatty acid metabolism genes [23] and interactions with Pdr1/3 to regulate the expression of drug efflux pumps involved in pleiotropic drug resistance [22, 35]. The KIX domain is also known to contribute to Gcn4 and Gal4 activation alongside other activator binding domains (Fig. 1A, ABD1-3 annotations) [20, 28, 36, 37].

Here we use ketoconazole tolerance as a phenotypic indicator of Pdr activity. Constructs lacking the KIX domain exhibited mild sensitivity to the drug, however a construct containing the KIX domain but lacking other ABDs was also mildly sensitive, suggesting the involvement of other Med15 regions in the Pdr response (Fig. 1B). In contrast, the presence or absence of the KIX domain alone (compare KIXΔ to *MED15*) appeared to have little detectable impact on Msn2 (heat, osmotic stress) or Gal4 (growth on galactose as sole carbon source) and Gcn4 (growth on SD media) phenotypes. The importance of the KIX domain in combination with ABD1-3 for Gal4 dependent activation could be seen in the KIX domain-containing *med15* 153-687Δ mutant, which exhibited better growth on galactose plates compared to the KIX-less *med15* 46-619Δ mutant.

In constructs lacking the KIX domain, the ABDs were important. For example, both KIX277-799Δ -*MED15* and KIXQ2Q3Δ-*MED15*, which lack the KIX domain but contain either a single (ABD1; KIX277-799Δ) or three ABDs (KIXQ2Q3Δ) grew well or very well on galactose (Fig. 1B), although both exhibited a modest reduction in expression of a UAS_G_-*lacZ* reporter relative to wild type (Fig. 1C). Similarly, growth on SD media, a reflection of Gcn4 activity, was poor in the 46-619Δ mutant as expected for a protein lacking the KIX as well as all ABD domains and best in the KIXQ2Q3Δ mutant which lacks the KIX domain but retains ABD1, ABD2 and ABD3 (Fig. 1B).

### Med15 Phosphorylation

In unstressed cells Med15 is phosphorylated, a modification proposed to dampen Med15 activation of stress genes. In contrast, Med15 is dephosphorylated in cells exposed to osmotic challenge [38]. There are 30 phosphorylated residues in Med15, all clustered around the junction of the central disordered domain (distal to Q3) with MAD (Fig. 1A) [38]. Of these, the phosphorylation of only 7 of these is affected by osmotic stress. In the D7P mutant, each of these seven dynamic phosphosites was changed to alanine, whereas in the D30P mutant all 30 phosphosites were alanine substituted [38]. We observed a growth defect in the D30P mutant at high salt concentrations but not D7P, as previously reported [38]. We found that these mutants also differed in response to acetic acid and ketoconazole. D30P exhibited wild type tolerance to acetic acid stress and ketoconazole, while D7P was more sensitive (Fig. 1B).

### The Central Q-rich and Intrinsically Disordered Region

We simplified the analyses of the central disordered domain and the Q1 tract by analyzing them in the absence of the KIX domain and the Q2/Q3 tracts. The KIXQ2Q3Δ-*MED15* construct, which lacks the KIX domain as well as the Q2 and Q3 glutamine tracts, nevertheless fully complemented all tested phenotypes except for ketoconazole and acetic acid tolerance when expressed in a *med15Δ* background (Fig. 1B) suggesting that the region around Q1 is more important than Q2 and Q3 for most KIX domain-independent Med15 functions. Interestingly, acetic acid tolerance, a likely function of the recently uncovered interaction between Med15 and Hap5 [39], was sensitive to the absence of Q2 and Q3 (Fig. 1B).

### The Q1 Tract

KIXQ2Q3Δ-*MED15* constructs containing a S288C (wild type strain background) size Q1 tract consisting of 12 glutamines (12Q) fully complemented several of the tested phenotypes including the utilization of galactose as a carbon source; sensitivity to ethanol; and sensitivity to NaCl (Fig. 1, Fig. 2A). However, a construct lacking the Q1 tract (0-Q1) neither fully complemented *med15Δ* defects on plates (Fig. 2A), nor UAS_G_-*lacZ* reporter activity (Fig 2C). One exception to the impact of removing the Q1 tract was that growth on minimal media (Fig. 2, SD +LKHU), which reflects Gcn4 activity, was not detectably affected.

**Figure 2.**
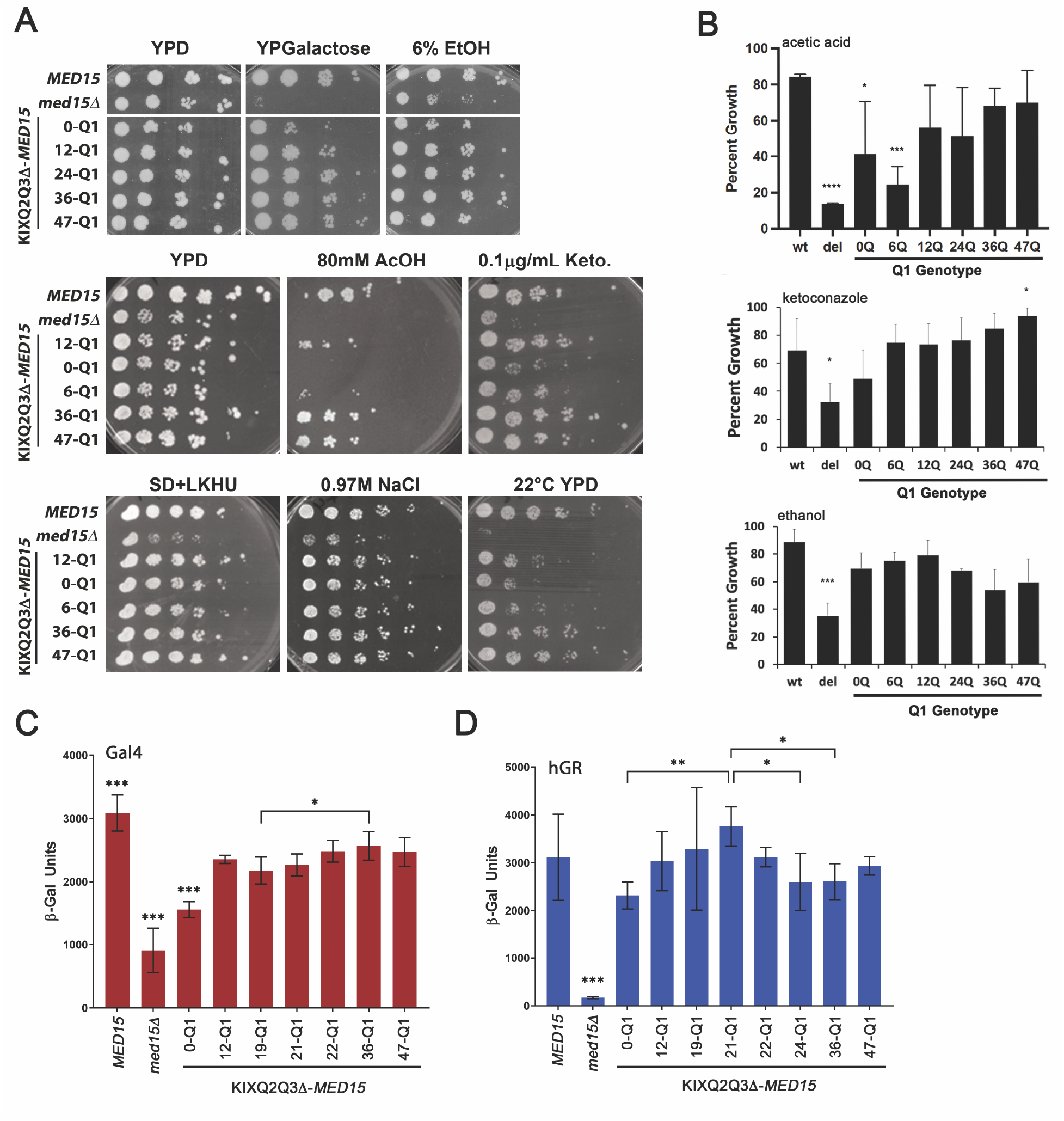
Poly-Q length variation at Q1 alters Med15 activity. **(A)** Log-phase cultures of the *med15Δ* strain (OY320) carrying a CEN plasmid with the indicated *MED15* or KIXQ2Q3Δ-*MED15* allele were serially diluted and spotted on YPD media with or without supplements and incubated at 30°C or room temperature (22°C) as specified. **(B)** Quantitative microtiter dish assay for growth in various media or in the presence of added chemicals. Each genotype was tested in biological triplicate. Plotted values are averages of the endpoint OD_600_ divided by growth in permissive conditions (media without chemical, 30°C). Drug concentration was chosen for an effect on the wild type *MED15* strain of no more than 50% reduction in growth and was 62.5 mM acetic acid, 6.25% ethanol and 5 μg/mL ketoconazole. Significance is indicated relative to growth of the wild type *MED15* strain. **(C-D)** β-galactosidase activity of a Gal4-dependent (UAS_G_-*lacZ*) reporter (C, red) as in Figure 1, or a human glucocorticoid receptor dependent (hGRτ1-*lacZ)* reporter (D, blue). All data points are the averages of at least 3 biological replicates (transformants) and error bars are the standard deviation of the mean. Significant differences were determined using ANOVA analysis with a Tukey post-hoc test, * p≤0.05.

To determine whether changes in the length of polyglutamine tracts, which are known to be naturally variable [21], modulate the activity of Med15, we tested a series of natural (12Q, 19Q, 21Q, 22Q) and synthetic (6Q, 24Q, 36Q, 47Q) Q1 tract lengths. Tract length effects at Q1 were context dependent. Plate phenotypes were examined and additional resolution achieved by quantifying growth in liquid media containing a stressor and comparing it to growth in media with no addition. In general, 0Q (and 6Q) inserts at Q1 afforded better tolerance than the *med15* deletion strain, but less than 12Q. In some cases, 36Q and 47Q constructs exceeded 12Q (Fig. 2B: acetic acid, ketoconazole) and in other cases 36Q and 47Q were less robust than 12Q (Fig. 2B: ethanol).

We also used an hGR reporter system [18] to examine the impact of tract length. The glutamine-rich domain of Med15 (aa 116-277) is similar in sequence to the glutamine-rich domains of human steroid receptor coactivators (SRCs) and thus the Q1R domain of Med15 in a construct equivalent to KIX277-799Δ fully activates a human glucocorticoid receptor (hGR) dependent report in yeast [18] as does the KIXQ2Q3Δ construct (Fig. 2D). In this context, a 0Q-Q1 tract reduced reporter activity and a Q1 length of 21 glutamines (Fig. 2D) exhibited a significant increase in hGR activity relative to longer tracts consisting of 24 or 36 glutamines. In contrast, changes to the Q1 tract length had only a minor impact on Med15 activation of a Gal4-dependent reporter (Fig. 2C).

Gcn4 is known to phase separate with a Med15 (aa 6-651) truncation encompassing 60% of the full-length protein [30] as part of a large transcriptionally active liquid droplet with phase separation dependent on Gcn4 residues essential for transactivation [30], indicating that liquid droplet formation is a key aspect of Gcn4-mediated activation. To determine if there is any impact of the Q1 tract on Med15-Gcn4 phase separation we first determined whether the Q1R domain could support phase separation on its own. This Med15 fragment is 157 residues and has an average disorder score over the entire region of 0.65 (IUPRED3 https://iupred.elte.hu/) with disorder peaking between amino acids 122 and 159 with an average score of 0.9. The region has an extreme glutamine/asparagine bias totaling 44.6% mixed with alanine (8.9%), leucine (8.9%) and proline (7.6%). We purified a full-length GFP tagged Gcn4 protein and various derivatives of mCherry tagged Med15 using nickel chromatography (Fig. S2) and used the proteins in LLPS assays. We found that mCh-Med15 proteins containing both the KIX domain and Q1R (KQ) as well as the Q1R only (Q1R) each coalesced with GFP-Gcn4 with moderate levels of PEG added as a crowding agent (Fig. 3A-B). The condensates did not form in the presence of 10% 1,6 hexanediol (Fig. 3C) which interferes with weak hydrophobic protein-protein interactions that are characteristic of liquid condensates.

**Figure 3.**
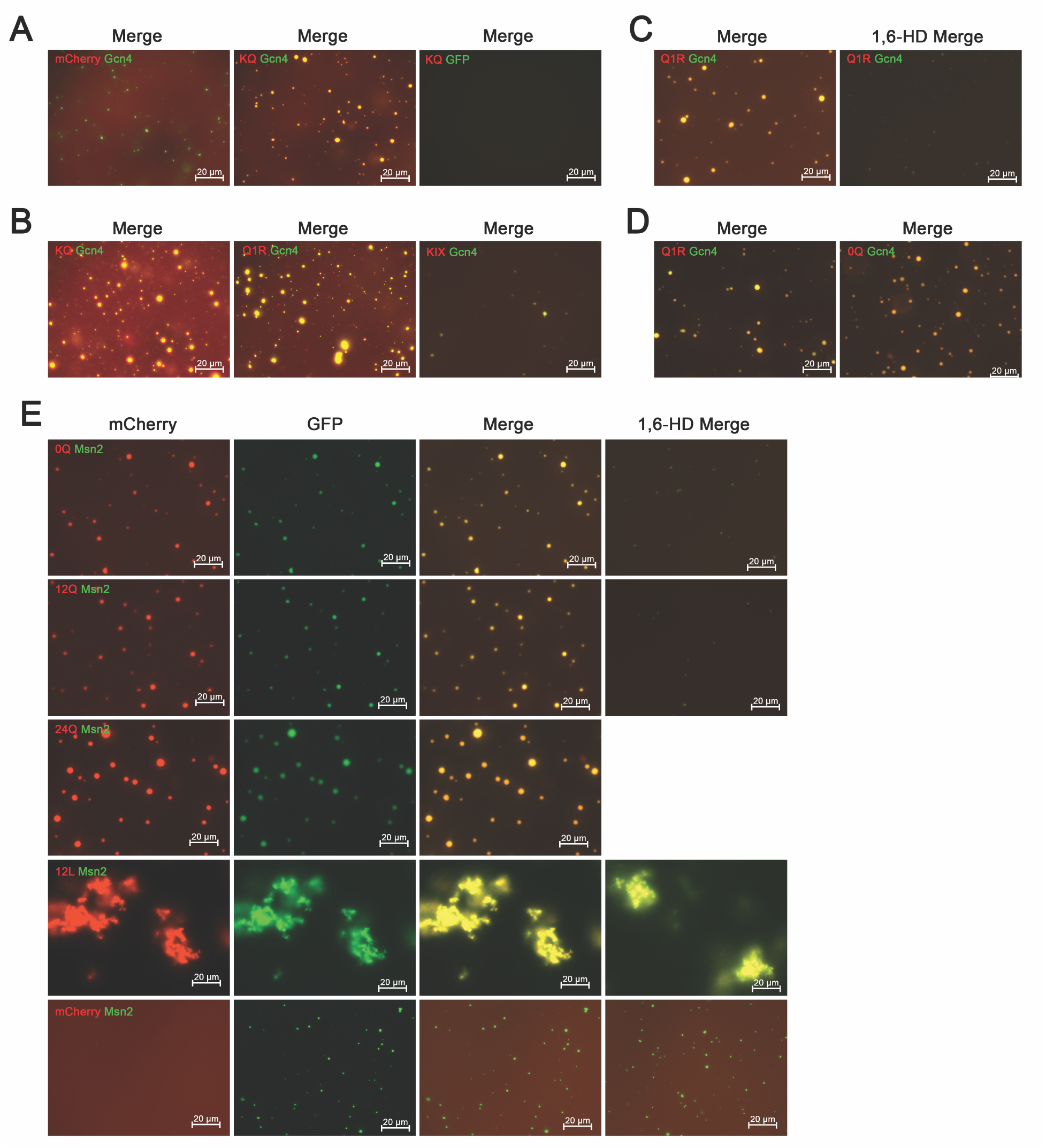
The Q1R fragment of Med15 forms liquid droplet like condensates with Gcn4 and Msn2_TAD_. **(A)** Representative microscope images of in vitro reactions consisting of 5uM Gcn4 and 20uM Med15 in the presence of 4% PEG and 125 mM NaCl. Images are the merge of the red and green channels. (**B)** The KIX-Q1R (KQ), Q1R and KIX (K) regions of Med15 were tested for droplet formation with Gcn4. (**C)** The Q1R-Gcn4 reaction was performed in the presence of 10% 1,6% 1,6-Hexandiol. **(D)** Comparison of Gcn4 aggregates formed in the presence of Med15 Q1R with and without the 12Q tract (Med15 12-Q1 Q1R and Med15 0-Q1 Q1R). **(E)** Representative microscope images of *in vitro* reactions consisting of 20 μM mCherry-tagged Med15 Q1R variants and 5 μM GFP-tagged Msn2 (aa 2-301), or the same concentration of mCherry in the presence of 6% PEG. 1,6 hexanediol was added to 10% in parallel 12Q and 0Q – Msn2 reactions and only the merged image is shown at the end of each of the first two rows. Likewise, 1,6 hexanediol was added to 6% in parallel 12L and mCherry reactions and only the merged image is shown at the end of each of the last two rows.

Having established that Q1R was sufficient for phase separation with Gcn4, we compared the phase separation characteristics of equivalent concentrations of purified Med15 Q1R and Med15 0Q-Q1R proteins in combination with low levels of Gcn4. No major effects of the Q1 tract on phase separation could be detected using microscopy (Fig. 3D).

We also examined phase separation of Med15 Q1R and 0Q-Q1R with the transactivation domain of Msn2 (aa 2-301). Microscopy was conducted to visualize the effects of the two Med15 Q1R variants at a single Med15 concentration (20 μM) in conjunction with 2.5 μM Msn2. This concentration of Msn2 formed small condensates in the presence of mCherry that did not require Med15. Added Med15 Q1R (12-Q1 and 24-Q1) coalesced with Msn2 condensates and increased their size. Added Med15 0-Q1 Q1R was also capable of forming condensates with Msn2. 10% 1,6 HD reduced condensate size and number in both the 0-Q1 and 12-Q1 reactions, (Fig. 3E). Interestingly, Med15 with a 12L-Q1 insert formed non-droplet aggregates that were resistant to 6% 1,6 HD.

Taken together, both macroscopic and microscopic assays suggest that while the Q1 tract is not essential for the Med15 activities tested, it is required for normal (wild type) levels of some Med15 activities, and different Med15 activities are differentially sensitive to the length of Q1.

### The Coiled-Coil Propensity of the Q1 Tract Influences Med15 Activity

The sequence requirements at the Q1 locus were investigated using a series of modified Q-tracts and non-Q insertions into the Q1 position of the KIXQ2Q3Δ construct. Since the region around the Q1 tract is predicted to form coiled-coil structure (Fig. and Fig. S3) that could be involved in protein-protein interactions, we tested a flexible glycine-rich spacer sequence (spacer: SPGSAGSAAGGA), predicted to have minimal coiled-coil propensity. The spacer construct was comparable to 12-Q1 under conditions in which the absence of the Q1 tract impaired Med15 activity (Fig. 4B, spacer). The Q1 locus was also replaced with a glutamine tract with dispersed proline residues (12PQ: PQQQPQQPQQQP and doubled, 24PQ), a sequence previously shown to disrupt coiled-coil structure [14] and that is predicted to reduce the coiled-coil structure of the adjacent region in Med15 (Fig. 4A and Fig. S3). The proline interrupted sequences also displayed activity comparable to 12Q-Q1 (Fig. 4B). In contrast, the introduction of leucine residues (15LQ; LQQQLQQLQQQLLLQ) previously shown to promote coiled-coil structure [14] did not complement the defects exhibited by the 0-Q1 construct (Fig. 4B top, most evident on galactose media). These results suggest that Med15 functionality does not require a coiled-coil prone sequence and that enhancing the coiled-coil propensity of the sequence at Q1 may be deleterious.

**Figure 4.**
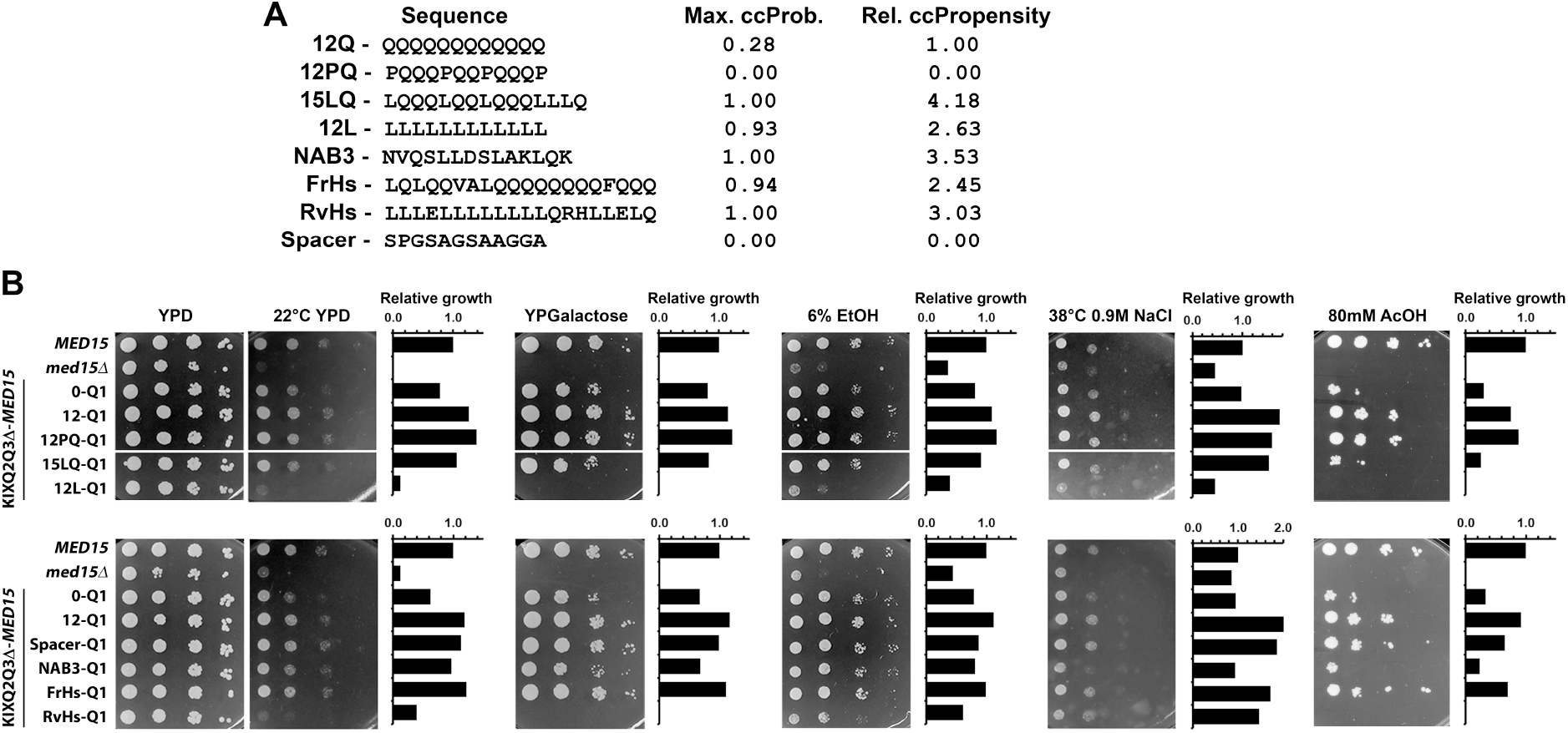
Q1 tract region coiled-coil structure perturbs Med15 activity. **(A)** Sequence and coiled coil (cc) propensity of non-Q inserts. Maximum ccProbability (Max ccProb.) corresponds to the peak coiled-coil formation probability within the Q1 region for each construct. Relative ccPropensity (Rel. ccPropensity) is the average ccProbability across the Q1 region peak relative to the 12Q construct prediction. **(B)** Log-phase cultures of the *med15Δ* strain (OY320) carrying a CEN plasmid with the indicated *MED15* allele were serially diluted and spotted on YPD media with or without supplements and incubated at 30°C or room temperature (22°C) as specified. Graphs provide a visual representation of growth for each row of each plate. Relative growth is a pixel-based growth measurement that incorporates both viability and colony size relative to the wild type *MED15*^+^ strain on each plate.

To further investigate the apparent deleterious effect of coil-coil promoting sequences, additional coiled-coil forming structures [5, 16] were introduced. The first was a C-terminal helix forming sequence from the *NAB3* gene (NVQSLLDSLAKLQK). Like 15LQ-Q1, NAB3 failed to fully complement the defects of the 0-Q1 construct (Fig. 4B: most evident on acetic acid containing media). One further test involved a 20 amino acid segment of the human Med15 Q-tract sequence (FrHs: LQLQQVALQQQQQQQQFQQQ) whose coiled-coil structure has been confirmed by circular dichroism [16]. The functionality of the human *MED15* Q-tract coding sequence was roughly comparable to 12-Q1 (Fig. 4B: FrHs-Q1). In contrast, the leucine rich sequence encoded by the reverse complement of that sequence (LLLELLLLLLLLQRHLLELQ) (RvHs-Q1; a periodically interrupted poly-L tract) and an uninterrupted poly-L tract (12L-Q1) exhibited only slightly more complementation in these assays than a *med15* deletion (Fig. 4B).

The results of these analyses suggest that Q1 inserts with weak or no predicted coiled-coil propensity (Fig. S3) are compatible with Med15 function, and that Q1 inserts that increase predicted coiled-coil propensity are less so.

The effect of length and composition of the Q1 tract on target gene expression was quantified by qRT-PCR in log-phase cultures (rich media, 30°C) of strains expressing wild type *MED15* or KIXQ2Q3Δ-*MED15* constructs with various Q1 genotypes. We chose to examine the effect of Med15 on basal expression rather than on induced expression for this analysis because of the major effect Med15 has on basal levels [40, 41]. We found that although induced levels were also affected, the extent of induction was not. The effect of *med15*Δ on basal expression of selected *MED15* target genes is shown in Fig. 5 and Fig. S4. We found that basal expression was dependent on the presence of the Q1 tract and that Q1 tract length variants gave distinctive patterns of expression. A Med15 derivative lacking the Q1 tract (Q1-0) was less active than Med15 with a Q1 tract of 12 (wild type) for genes such as *AHP1* (activation) and *MET10* (repression) (Fig. 5A). Although the basal expression of *MET10* and *AHP1* was dependent on the Q1 tract, it was independent of tract length. In contrast, *GLK1* was sensitive to tract length, with significantly lower expression in Med15 variants with a Q1 tract length of 36 or more. The *SSA1* and *HSP12* expression patterns showed a similar trend with the lowest expression in Q1-36 and Q1-47 Med15 variants (Fig. 5A).

**Figure 5.**
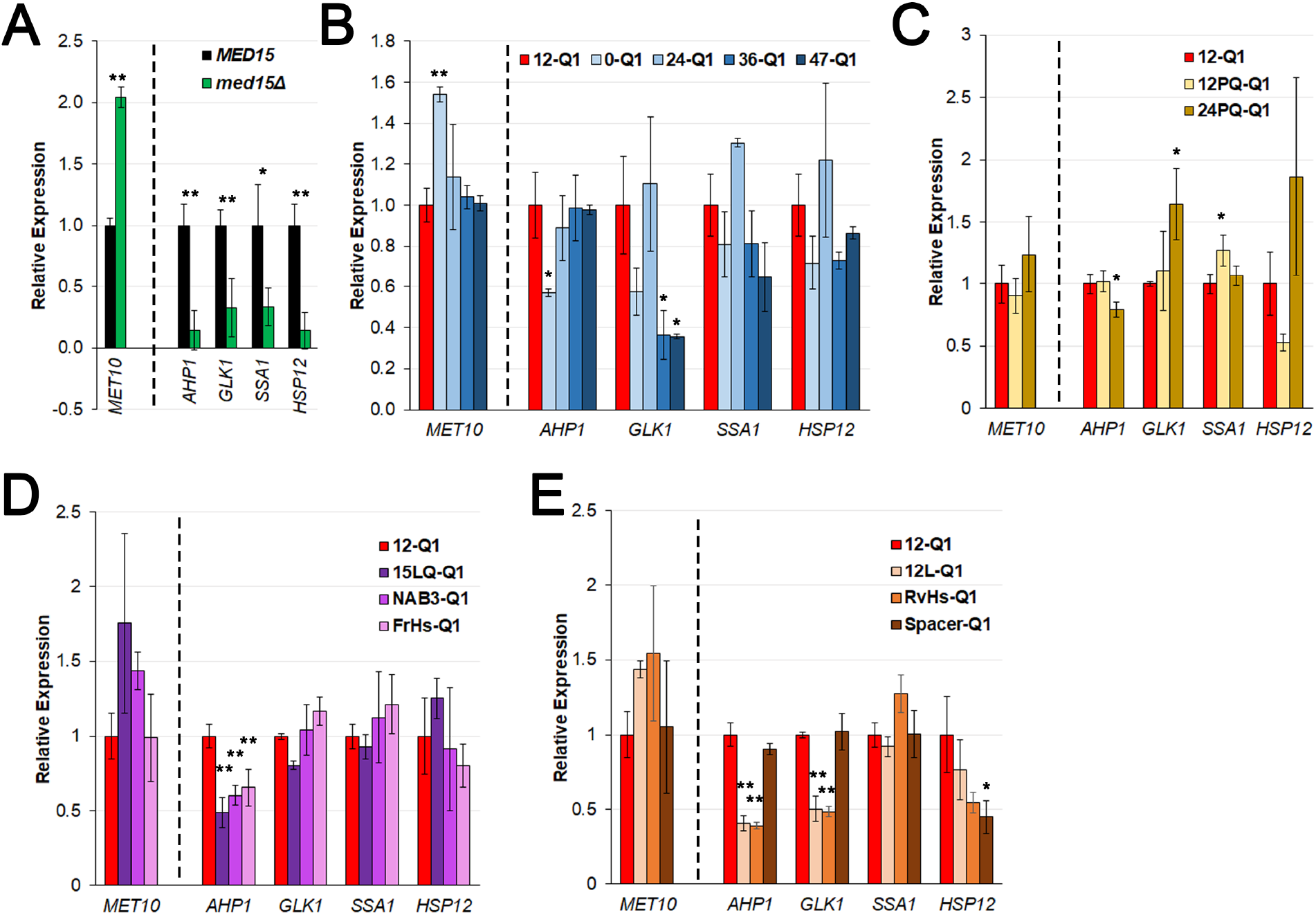
Basal Gcn4 and Msn2 dependent gene expression is modulated by the presence and length of the Q1 tract in Med15. Relative expression of target genes for log-phase cultures of the *med15Δ* strain (OY320) carrying CEN plasmids with specified KIXQ2Q3Δ-*MED15* Q1 variant constructs in YPD grown at 30°C. Expression values for 12Q-Q1 (red) are set to 1. **(A)** *MED15* dependence of five representative target genes under basal (uninduced) conditions. **(B)** Q1 length variants; **(C)** Q1 inserts with interspersed proline variants designed to disrupt possible coiled-coil structures; **(D)** Q1 inserts with predicted or established coiled-coil structure **(E)** Non-glutamine Q1 inserts. For each strain the target gene expression was normalized to *ALG9* and averaged over 3 biological replicates (transformants). Significance was determined for each group of sequences relative to 12Q-Q1 using ANOVA analysis with a Tukey post-hoc, * p≤0.05, ** p≤0.01.

We also examined gene expression in the constructs with Q1 substitutions (Fig. 5B). For *AHP1* and *MET10,* genes whose expression was most consistently influenced by the presence of the Q1 tract, the introduction of proline residues into a 12-residue glutamine-rich tract did not significantly alter expression while the introduction of proline residues into a 24-residue glutamine-rich tract was somewhat deleterious. However, this effect was not evident in *GLK1*, *SSA1* and *HSP12* where the presence of prolines in a glutamine-rich tract of 12 or 24 appeared to increase expression (Fig. 5B).

Coil-promoting inserts such as FrHs-Q1, NAB3-Q1, 15LQ-Q1, and 12L-Q1 adversely affected *AHP1* and *MET10* expression, but other genes were not affected (Fig. 5C). Finally, leucine rich inserts such as 12L and RvHs Q1 were deleterious for *MET10* and *AHP1* expression as well as *GLK1,* while *SSA1* was unaffected. A non-Q spacer insert of 12 residues was neutral for all genes except *HSP12*, where it had an adverse effect (Fig. 5D).

Overall, Q1 tract composition differentially affected the expression of this panel of genes, consistent with the phenotype studies in Fig. 4.

### Med15 interacts with Msn2 via the KIX domain or the Q1R region

To determine whether Q1-dependent changes in the activity of Med15 reflect altered interactions with transcription factors, we examined Med15-Msn2 interactions using a split-ubiquitin two hybrid assay with a Ura3 reporter [42, 43]. Msn2 amino acids 1-271 corresponding to the transcriptional activation domain (TAD) were fused to the C terminus of ubiquitin and an N-end rule sensitive derivative of the *URA3* gene (R*URA3*), and different segments of Med15 were fused to the N terminus of ubiquitin (Fig. 6A). The presence of interacting N-Ub and C-Ub fusions cause strains to become Ura^-^ due to the ubiquitin-mediated degradation of the Ura3 protein. The Ura^-^ phenotype was tested in two ways: (1) with synthetic media containing lysine (SD+K) or synthetic complete media lacking uracil (SC-HLMU) on which strains harboring plasmids expressing fragments of Msn2 and Med15 that do interact would fail to grow, and (2) on media containing 5-fluoroorotic acid (5-FOA), an inhibitor of the Ura3 enzyme, on which only strains expressing interacting Med15 and Msn2 peptides would survive. Both the KIX domain alone (NUb-KIX; (aa 1-118)) and the Q1R fragment alone (NUb-Q1R-12Q; (aa 120-277)) as well as the entire region (NUb-KQ; (1-277)) were positive (Ura^-^ and 5-FOA^R^) for an interaction with the transactivation domain of Msn2 (Fig. 6B).

**Figure 6.**
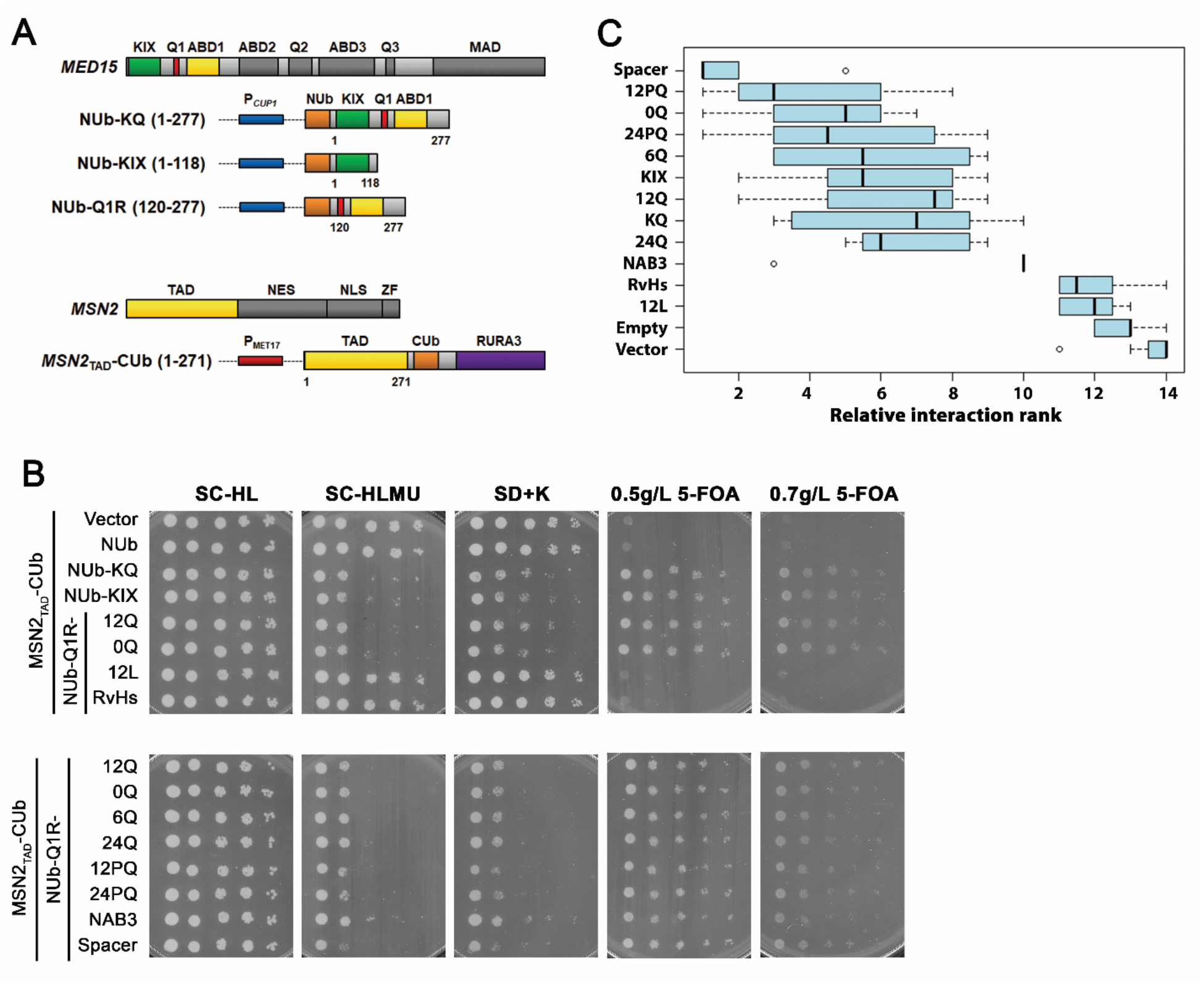
Med15 interacts with Msn2 through the KIX domain and Q1R. **(A)** Split ubiquitin constructs. Wild type yeast (BY4742) transformed with plasmids expressing N-terminal Ubiquitin fragment (NUb) fused to fragments of *MED15* behind a copper inducible *CUP1* promoter and C-terminal Ubiquitin fragment (CUb) fused to a mutated *URA3* gene coding for an N-terminal arginine (R*URA3*) and the transcriptional-activation domain of MSN2 (*MSN2*_TAD_) behind a methionine repressible *MET17* promoter. **(B)** Log-phase cultures in minimal media with leucine, lysine, and uracil (lacking methionine) were exposed to 0.05 or 0.2mM copper for 1 hour and then serially diluted and spotted on media lacking uracil or containing 5-FOA. Plates corresponding to strains induced with 0.2mM copper are shown. **(C)** Relative spot size formation quantified per plate as in Figure 3 at a single time point for plates lacking uracil or containing 5-FOA corresponding to strains induced with 0.1 or 0.2mM copper (8 total plates per strain). The relative growth rank was determined for each plate across all 14 tested constructs. A rank of 1 corresponds to the worst growth on plates lacking uracil and best growth on plates containing 5-FOA.

### Substitutions at Q1R affect the Msn2 interaction

Consistent with the phenotypic analysis shown in Fig. 4, an NUb-Q1R with *NAB3* sequence at Q1 was slightly less interaction positive (more Ura^+^) than Q1R itself (best seen on SC-HLMU in Fig. 6B), while a Q1R with RvHs or 12L at Q1 only weakly interacted with Msn2 (best seen on 5-FOA in Fig. 6B) despite being positive for protein expression (Fig. S5).

Interactions were ranked based on pixel density from high (least growth on plates without uracil or most growth on plates with 5-FOA) to low (most growth on plates without uracil or least growth on plates with 5-FOA) and a composite interaction rank based on growth on all plate types split the tested constructs into 3 groups (Fig. 6C). The poorest interactors, RvHs and 12L, ranked alongside the negative controls. The NAB3-Q1 construct was the only tested representative of the coiled-coil-promoting constructs, and it was in a group of its own as the weakest interactor except for the non-functional constructs. Interestingly, there was very little variability in the interaction score from plate to plate, so the NAB interaction rank appears very discrete in Fig. 6C (*NAB3*). The strongest interactor was the Spacer construct, which slightly outperforms the remaining tested constructs. Hence the Q1 substitution phenotypes reflect the ability of Med15 to interact with Msn2.

Interestingly, despite a robust interaction between the Msn2 transactivation domain and the Med15 KIX domain detected using split-ubiquitin two hybrid analyses, Med15 did not contribute appreciably to the small condensates formed by Msn2 alone. A low level of the Med15 KIX domain joined small Msn2 condensates (Fig. 7) but there was no evidence that the presence of Med15 KIX facilitated or stimulated Msn2 phase separation as the condensates were no larger or more abundant than those formed with an equivalent concentration of mCherry with no associated Med15. We surmise that the interaction between Msn2 and Med15 has two components, one that is driven by multivalency and one that is more likely driven by specific residue-based electrostatic interactions. The absence of any contribution of the KIX domain is mirrored in the condensate reactions between Med15 and Gcn4 (Fig. 3B).

**Figure 7.**
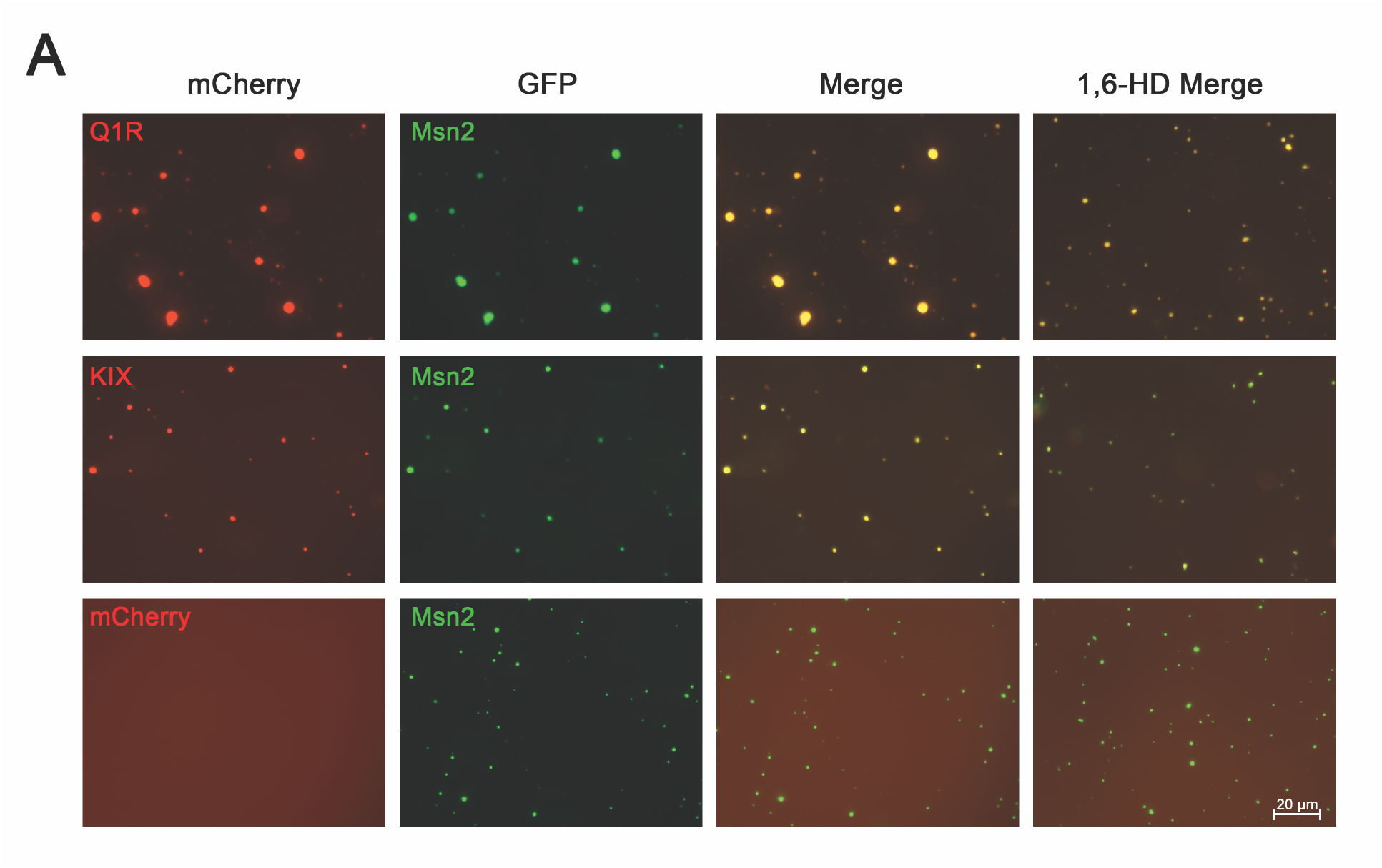
The KIX domain of Med15 does not undergo efficient phase separation with Msn2. Representative microscope images of 20μM mCherry tagged Med15 fragments (Q1R or KIX) or 20 μM mCherry with no associated Med15 together with 5 μM GFP tagged Msn2 in the presence of 6% PEG-8000 in the presence or absence of 6% 1,6 hexanediol.

### Med15 – Med15 interactions

To examine potential interactions within Med15 as suggested by the effect of distal regions of the protein on KIX-dependent activities, a series of NUb fusions with different segments of Med15 were tested with a CUb-KIX construct. There was a modest interaction between the KIX domain (aa 1-118) and Q1R (aa 120-277) (Fig. 8B, row 3). To probe the importance of the Q1 tract in the interaction with the KIX domain, we examined the effect of several Q1 length and composition variants on the KIX-Q1R interaction. Most Q1 substitutions had little to no effect on the interaction with the KIX domain, however, the leucine rich insertion, 12L (Fig. 8C) was negative for the interaction with the KIX domain, suggesting that the failure of rich Q1R substitutions to interact with Msn2 may reflect a non-permissive intramolecular interaction within the Med15 protein.

**Figure 8.**
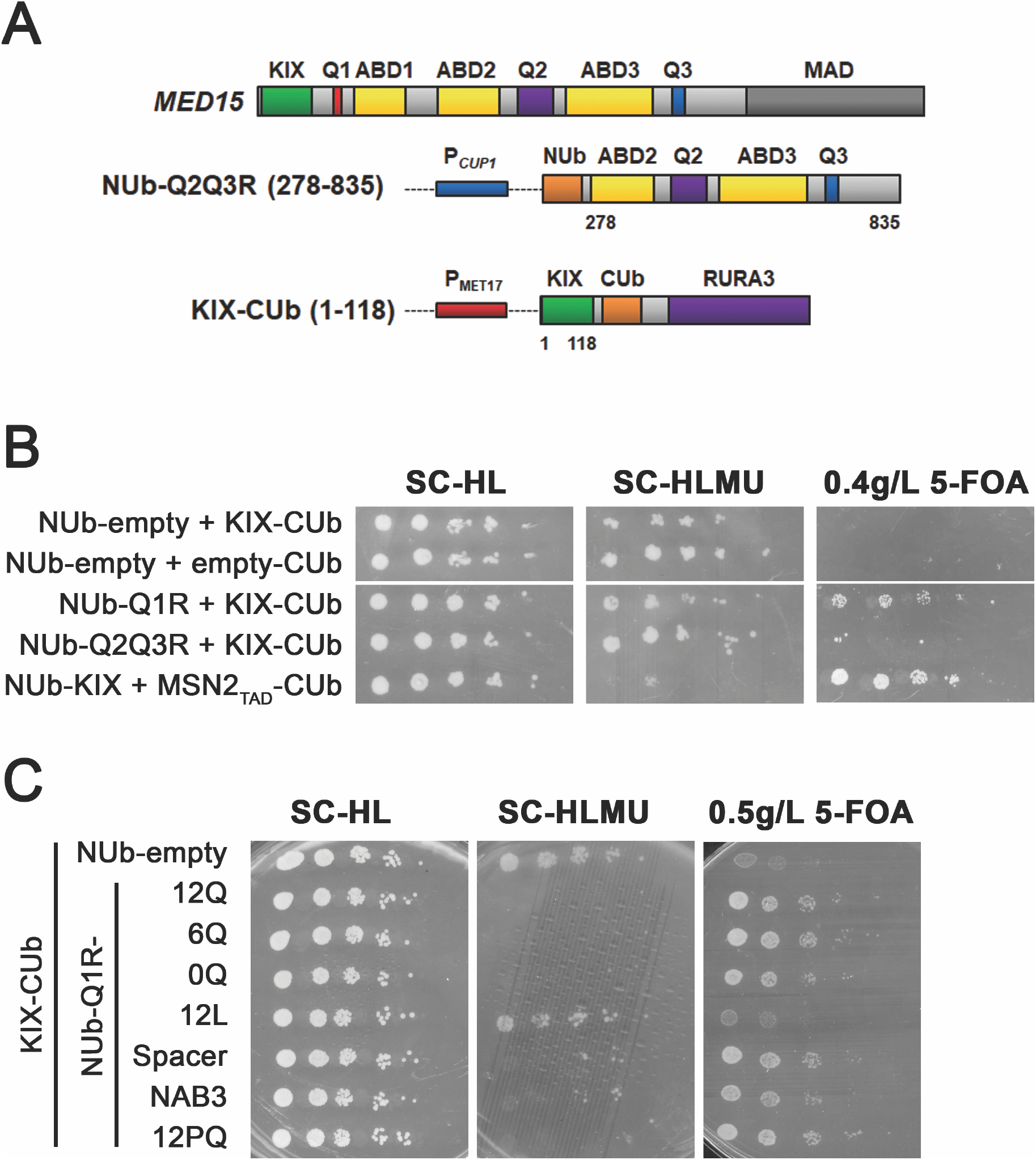
Interactions between Med15 sequences. **(A)** Cartoon depicting split ubiquitin constructs. **(B)** Log phase cultures of wild type strains carrying both CUb and NUb plasmids were exposed to 0.05 mM copper (and in some cases 0.02 mM methionine) for 1 hour and then serially diluted and spotted on media lacking uracil (SC-HLMU) or containing 5-FOA. (**C)** Substitutions of the 12Q region in Q1R were tested for effects on the interaction with the KIX domain.

## Discussion

In this study we used internal deletions as well as synthetic Med15 derivatives to specifically interrogate the role of the N-terminal most polyglutamine tract (Q1) adjacent to Activator Binding Domain 1 (ABD1). All *MED15* variants tested were built in the context of the *MED15* promoter and 3’ UTR sequences within a *LEU2*-marked centromeric plasmid. We used qRT-PCR and Western analysis to confirm that the expression of each construct was at least as robust as constructs that exhibited completely wild type functionality (Fig. S1). In earlier studies, Jedidi *et al*., [36] measured protein levels of many internal deletions of a myc-tagged *MED15* gene, and found that constructs lacking amino acids 169-202 or 500-543 were not expressed well. Two of the constructs depicted in Fig. 1, Δ153-687 and Δ46-619 lack these sequences, however in Western analysis of the N terminally 1xHA-tagged Δ46-619 construct, the protein was in fact expressed (Fig. S1D), thus ruling out the possibility that the phenotypes in Fig. 1 are confounded by reduced levels of Med15 protein. The difference in protein abundance and stability in our study and the Jedidi *et al*. [36] study may be due to the difference in tag or, more likely, the difference in the full complement of retained/absent sequences.

Using these constructs, we systematically analyzed the impact of different regions of the Med15 protein and found that the KIX domain and Q1R region of the protein are key determinants of Med15 activity (Fig. 1) although more distal parts of the protein, including Q2, Q3 and phosphorylated residues that span the boundary with the Mediator Association Domain appeared to influence the functionality of these more N-terminal regions suggesting that post-translational modifications or intramolecular interactions may also play a role.

Additional insight into the role of the Q1R region of Med15 was achieved by varying the length and composition of the Q1 tract in constructs lacking the KIX domain as well as the Q2 and Q3 tracts. In these analyses, we probed the activity of the Med15 protein using both phenotypic and target gene expression assays for the output of different Med15-dependent transcription factors. We anticipated that different Med15-TF interactions would depend on distinct regions of Med15 as previously shown [36], and that there might also be a TF-specific response to the presence/absence and content of the Q1 motif since TF interaction and activation domains fall into different classes (fuzzy, well structured, acidic etc.). The Med15 interaction with Gcn4 was previously shown to require dispersed ABDs in what has been described as a fuzzy interface [28, 44, 45] and could conceivably be independent of Q1, whereas Msn2, with its well-structured Med15 interaction motif [43], might be expected to behave differently.

The results of our analyses are summarized in Table 1. We found that activity of most tested TFs was diminished if the Q1 tract was deleted. We further showed that several TFs except for Gcn4 and Gal4 were affected by the length of the Q1 tract, with the precise effect being context dependent. Finally, we found that the amino acid composition and/or secondary structure was a factor in the activity of Med15-dependent TFs. We observed that Q1 substitutions predicted to increase coiled-coil propensity (Fig. S3) diminished TF activity while Q1 substitutions predicted to interfere with coiled-coil propensity had no effect on TF activity (Fig. 4, 5), suggesting that flexibility of the sequence is an important feature. Below we discuss each of these observations in more detail.

**Table 1.** Relationship of Med15 Q1 Genotype to TF Activity.

| Phenotype | TF | Target gene /<br>reporters<br>examined | Sensitivity to Q1<br>(phenotype / gene(s)) |  |  |
| --- | --- | --- | --- | --- | --- |
|  |  |  | Presence<sup>1</sup><br>(Fig. 2,3) | Length<sup>2</sup><br>(Fig. 2,4) | Composition<sup>3</sup><br>(Fig. 3,4) |
| Acetic Acid | Hap5 | Plate assay | Yes | Yes | Yes |
| NaCl @ 38°C;<br>Ethanol | Msn2;<br>Hsf1 | Plate assays | Yes | Yes | Yes |
|  |  | GLK1 | Yes | Yes | Yes |
|  |  | AHP1 | Yes | No | Yes |
|  |  | HSP12 | Yes | Yes | Yes |
|  |  | SSA1 | Yes | Yes | Yes |
| Ketoconazole | Pdr1 | Plate assay | Yes | Yes | No*<sup>4</sup> |
| Galactose | Gal4 | Plate assay | Yes | No | Yes |
|  |  | UAS<sub>G</sub> reporter | Yes | No | NT<sup>5</sup> |
| Amino Acid<br>Limitation | Gcn4 | Plate assay | No | No | No |
|  |  | MET10 | Yes | No | No |
|  |  | ARG3 | No<sup>6</sup> | Yes* | NT |
| Suboptimal<br>Temperature<br>(22°C) | ?<sup>7</sup> | Plate assay | Yes | Yes | Yes |
| Glucocorticoid<br>Response | GR | hGR reporter | Yes | Yes | NT |

<sup>1</sup>Growth or expression altered by the removal of Q1
<sup>2</sup>Growth or expression altered by changes in the length of Q1
<sup>3</sup>Growth or expression altered by changes in the composition of Q1
<sup>4</sup>Data not shown\*
<sup>5</sup>Not tested
<sup>6</sup>Data in supplement
<sup>7</sup>Target transcription factor unknown

### The KIX-dependent Med15 interaction with Hap5 is affected by additional Med15 sequences

*HAP5* is essential for the DNA-binding activity of the HAP complex, a tripartite CCAT-binding factor, which regulates respiratory gene expression. Hap5 mutants are also deficient in growth on 0.3% (vol/vol; ∼50 mM) acetic acid [39]. The Hap5 interaction with Med15 is KIX-dependent as shown by standard two-hybrid assays. Mutations in the KIX domain result in Hap5-dependent phenotypes including sterol homeostasis and acid tolerance [39].

The acetic acid tolerance phenotype was among the most revealing of the *med15* mutant phenotypes we examined. We observed acetic acid sensitivity stemming from the absence of Q2 and Q3, and in the D7P mutant which eliminates the potential for phosphorylation of C terminal sequences required to prevent activation under non-stress conditions and which become dephosphorylated upon osmotic stress (Fig. 1) [38]. Acetic acid tolerance was also affected by the absence, length, and composition of the Q1 tract (Fig. 2, Fig. 4). Taken together the Med15 sequences required for acetic acid tolerance were consistent with either dispersed activator binding sites for the Hap5 acetic acid responsive TF [39] or a requirement for intramolecular interactions between different regions of the Med15 protein. For example, we hypothesize that an interaction between the phosphorylated region located near the C-terminal mediator association domain and N-terminal sequences like the KIX domain or Q1 tract could squelch Med15 activity by preventing association of the subunit with the remainder of the complex.

Msn2 TF activity is mediated by the KIX and the Q1R regions of Med15 and is sensitive to Q1 length variants and substitutions.

A miniaturized version of Med15 consisting of aa 116-277 (ABD1, Q1) and the Mediator Association Domain (KIX277-799Δ-*MED15*) [18] fully complemented some of the stress related defects of the *med15*Δ strain. The activity of this construct is consistent with, and further refines, previous work in which the N terminal 351 amino acids were found to be sufficient for interactions between Med15 and the stress responsive transcription factor Msn2 [43]. We confirmed that Msn2-related activities of Med15 are encoded by the region of Med15 containing the Q1 tract and ABD1 (aa 116-277) but found that the KIX domain alone could also mediate an interaction with Msn2 (Fig. 6), which was not previously appreciated. The fact that there was no additivity between Q1R-based and KIX-based interactions with Msn2 in functional assays (Fig. 1) or in the split ubiquitin two-hybrid interaction assays (Fig. 6) together with the absence of condensate formation between Msn2 and the KIX domain may reflect the presence of two types of Msn2-Med15 interactions that are mutually exclusive and potentially redundant.

### The presence and length of the Q1 tract affect Gcn4 and Msn2 activation

The effect of Q1 tract length on growth and stress response phenotypes and on expression of reporters and target genes was context dependent. Basal (Fig. 5) and induced (Fig. S3C) expression of individual Msn2-dependent (e.g., *AHP1*) and Gcn4 dependent (e.g., *MET10*) genes was influenced by the Q1 tract. In all instances TF activity was reduced in the absence of the Q1 tract. Note that Med15 is a negative regulator of *MET10* expression so *MET10* expression increases when Med15 activity decreases [21]. This suggests a general role for the Q1 tract in modulating Med15 dependent regulation of TFs, potentially in providing structural flexibility that facilitates TFs accessibility to residues in Med15 that participate in the interaction, although other mechanisms haven’t been ruled out.

Whether the length of the Q1 tract influenced gene expression depended on the transcription factor. Gal4-dependent reporter gene expression as well as the expression of Gcn4 regulated *MET10* were largely tract-length insensitive (Fig. 2C and Fig. 5A). The insensitivity of Gcn4 and Gal4 interactions with Med15 to Q1 tract length suggests that “fuzzy” interactions may require a threshold number of residues but are otherwise independent of length.

In contrast, expression of an hGR-dependent reporter peaked at 21Q and was reduced at 36Q (Fig. 2D). Similar expression patterns were seen for Msn2/Hsf1 targets including *HSP12*, *SSA1* and *GLK1* where expression peaked at 12-24Q and was reduced at lengths > 36Q (Fig. 5A).

With respect to Q1 tract composition, there was general agreement in gene expression patterns (Fig. 5B-D) and Med15 phenotypes (Fig. 4). The non-functional RvHs and 12L Q1 inserts exhibited the most extreme changes in gene expression, followed by the partially functional 15LQ and *NAB3* Q1 inserts. Med15 Q1 substitutions affected the interaction with the Msn2 transcription factor. The Msn2:Med15 interaction was most affected by Q1 substitutions like 12L and RvHs that were the least functional and was somewhat affected by the NAB3 insert which exhibited some reduction in functionality (Fig. 4, Fig. 6B, C). Specifically, the helical nature of the NAB3 insert sequence may promote a structure that alters the availability of adjacent residues needed for the Msn2 interaction. Hence, overall, we conclude that the nature of the Q1 sequence has an impact on the interaction with the Msn2 transcription factor, while the length of the glutamine tract at Q1 may impact Med15 function differently.

It is worth noting that gene expression is not necessarily a direct readout of the interaction strength between any single pair of proteins. The upstream regulatory regions of the genes analyzed in our studies contain a complex network of binding sites for positive and negative regulators that likely interact in uncharacterized ways, making it difficult to predict whether a strong or weak interaction between one transcription factor and one Mediator Complex subunit will lead to normal levels of gene expression. Hence, while it may be surprising, it is not incongruous that the Q1 inserts (Spacer and 12PQ) conferring the strongest Msn2 interaction (Fig. 6C) lead to reductions in the expression of the Msn2 target gene, *HSP12* (Fig. 5B). An additional layer of complexity is introduced because the interaction between Med15 and Msn2 not only recruits RNA Pol II to the target gene, but also brings Msn2 into proximity of the CDK module of Mediator that subsequently promotes the phosphorylation and degradation of Msn2 [31].

### The role of glutamine bias and coiled-coil propensity in Med15 Function

The extreme glutamine bias in yeast Med15 is conserved in other fungi as well as in animal orthologs [21]. The conservation of glutamine bias suggests a possible mechanistic basis for glutamine overrepresentation. Mechanistically, poly-Q tracts could influence activity in various ways: by affecting protein-protein interactions, either directly or indirectly; by providing disorder to allow a larger set of transient interactions; by providing necessary spacing between functional domains; or by providing flexibility to the protein. We found that the flexible spacer sequence (SPGSAGSAAGGA) worked as well if not better that the native 12Q sequence (Fig. 4). These results suggest that flexibility is more important than the sequence and that glutamine residues are not themselves required for creating or stabilizing a protein-protein interface. The fact that residues at Q1 were not functionally constrained to be glutamine implies that the Q1 tract is not itself an interaction motif participating directly in protein-protein interactions. This conclusion is supported by our observation that removal of the Q1 tract reduced but did not eliminate activity (Fig. 2). Furthermore, no previous studies have attributed protein interactions to the Q1 tract in Med15. In the context of the hGR reporter, two residues, Q198 and V199, downstream of the Q1 tract were found to be critical for the interaction with hGR τ fragment [18]. Gcn4, and more recently, Gal4, have been shown to interact with the ABDs of Med15 [44, 45]. While the Q1 tract is adjacent to ABD1, it is not a required component of the ABD1 interaction surface.

Glutamine rich sequences in Med15 and other proteins have been reported to confer functionally important coiled-coil structure [5, 14, 16]. However, we found that the presence of periodic proline residues known to perturb coiled-coil structures in circular dichroism studies of Med15-related peptides [14] did not impair activity. This observation is consistent with the naturally occurring proline interruptions in the glutamine-rich regions in functional animal orthologs of Med15 [21]. In contrast, we found that the insertion of sequences with established coiled-coil propensity such as the coiled-coil forming Q1 adjacent sequence in human Med15 [16] and especially the glutamine-adjacent region from the C-terminus of the Nab3 protein [5] dampen Med15 activity (Fig. 4), suggesting that torsional flexibility of the Q1 region may be important.

Interestingly, two Q1 substitutions were not tolerated in phenotypic assays (Fig. 4) and were interaction negative with Msn2 (Fig. 6). One was the 12L (inverted glutamine, CTG) sequence, and the other, the inverted Human Med15 coil (RvHs), which is also very leucine rich (Fig. 4). The basis for the deleterious effect of leucine-rich sequences is unclear, however, we found that leucine-rich Q1 tracts also interfered with an intramolecular interaction between the Q1R and the KIX domains (Fig. 8C). Finally, we observed that a 12L Q1R protein was negative for phase separation with both Gcn4 (data not shown) and Msn2 (Fig 3E). Interestingly both Gcn4 and Msn2 proteins formed geometric co-aggregates with the 12L version of Med15 suggesting that the proteins do interact *in vitro* but that the interaction is non-productive in vivo. Furthermore, we observed that the 12L insertion at Q1 position of the KIX277-799Δ-Med15 construct had a dominant negative effect when introduced into a wild type *MED15*^+^ strain (data not shown). These observations could reflect the disruption of a functionally important regulatory interaction. However, the possibility of off-target effects cannot be definitively ruled out as a cause of reduced TF interaction and diminished functionality of this Med15 variant.

### The role of glutamine bias in Med15 coalescence with liquid droplet condensates formed by TFs

In *in vitro* condensate reactions consisting of Gcn4 (full length) or Msn2 (TAD, aa 2-301) and Med15 Q1R or Med15 0Q-Q1R, we observed that the Q1 tract did not make major contributions to the ability of Med15 (Fig. 3B) to coalesce with the transcription factor. These observations are supported by turbidity measurements in which low levels of Gcn4 or Msn2 are mixed with increasing concentrations of Med15 and the turbidity due to aggregation (340 nm) plotted following a 1,6 HD correction of each data point to remove contributions from non-liquid type condensates (Fig. S6). This rules out one additional mechanism by which the Q1 tract might influence Med15 function.

### The role of disorder and spacing at Q1 of Med15

The possibility that glutamine enrichment, including the poly-Q tracts, found throughout the midsection of Med15 simply promote disorder remains plausible. By promoting disorder, poly-Q tracts could increase the number of structural conformations Med15 can assume as well as increase the number of potential compatible interactors. However, a role for the Q1 tract as a spacer, hinge [7] or conformational modulator [46] cannot be ruled out. In studies of the Huntingtin protein, the expansion-prone poly-Q tract in exon 1 appears to allow adjacent domains within a protein to interact [7]. Both the removal and extensive expansion of the poly-Q tract of the Huntingtin (Htt) protein disrupts the proper hinge function with a shorter Huntingtin Q tract of 6 glutamines being partially defective [7]. In separate studies, the polyglutamine expansion within Htt exon 1 was shown to lead to increased alpha helical structure affecting the curvature of the protein and altering intramolecular interactions with the remainder of the protein [46].

The shortest Q1 tract we identified in *MED15* from sequenced *S. cerevisiae* genomes is 10 glutamine residues [21]; Q1 tracts shorter than 10 do not occur frequently in natural populations. Although complete removal of the Q1 tract (Q1=0) does not lead to complete loss of Med15 function, the reduction in Med15 functionality is clear in every assay. We speculate that a compromised Med15 of this sort may be maladaptive and would be selected against in nature. Definitive tests of a hinge role for the Q1 tract of Med15 are potentially confounded by the participation of the Med15 protein in weak multivalent interactions in the presence of TFs like Gcn4 and Msn2 in the context of phase-separated liquid droplets. However, in testing for intramolecular interactions (Fig. 8) between sequences on either side of the Q1 tract, we found that the KIX domain is positive for interactions with Q1R. This type of interaction could be a reflection of novel mechanisms for regulating Med15 activity.

Our working hypothesis for the role of polyglutamine tract Q1 and other Med15 sequence features in modulating intra-and inter-molecular interactions of Med15 is depicted in Fig. 9. Various types of Med15 interactions are shown: (1) represents the interaction of Med15 with the remainder of the Mediator Complex. This interaction is primarily dependent on a specific motif in the MAD region of Med15 [47], and likely independent of Q tracts. (2) represents Med15 interactions with activator binding domains (ABDs) via fuzzy (weak multivalent) (a) or directed (b) associations. (3) points to the role of poly-glutamine tracts in providing flexibility and spacing between ABDs which influences binding and levels of transcriptional activation. And (4) depicts intramolecular interactions between regions of Med15. These interactions may serve to compete with TF interactors thus limiting the intermolecular interactions and consequently limiting the extent of transcriptional activation. Alternatively, these intramolecular interactions may serve to prevent aberrant interactions between Med15 and unintentional targets by obscuring interaction surfaces.

**Figure 9.**
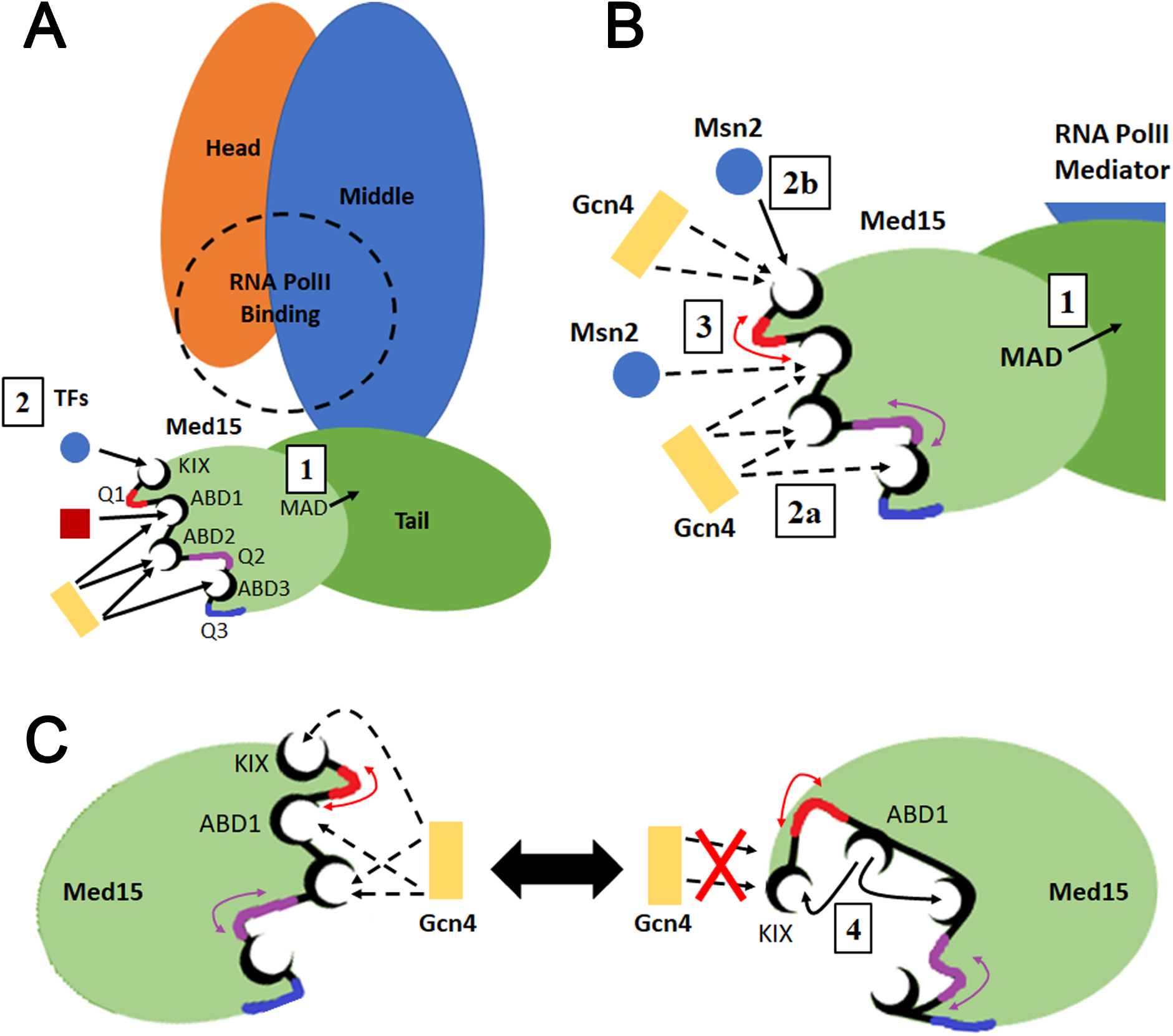
Cartoon model depicting different types of Med15 interactions. (A) View of Med15 in a pre-initiation complex. (B) Zoomed in view of a TF binding interface of Med15 with Gcn4 and Msn2 shown as examples. (C) A depiction of the alternate states of Med15 either forming intermolecular interactions with a TF (Gcn4 shown as an example) or forming intramolecular interactions. Specific types of interactions are numbered: (1) Integration into Mediator likely independent of Q-tracts and dependent primarily on MAD. (2) Different TFs interact with MED15 ABDs through fuzzy (a, dashed line) or directed (b, solid line) interactions. (3) Poly-Q tracts provide flexibility and spacing between ABDs which influences interactions and transcriptional activation. (4) Intramolecular interactions within Med15 that compete with or are displaced by TF intermolecular interactions.

The potential presence of two different types of Med15 interactions, conformationally flexible weak multivalent interactions (leading to phase separation) such as Med15:Gcn4 and more stoichiometric interactions such as the one we postulate between the Med15 KIX domain and Msn2 or Gcn4 would not be unusual. Fuzzy interactions are ubiquitous in stoichiometric protein complexes [48-50]. The precise structural contributions of the potential for fuzzy complexes in Med15 are yet to be determined. For example, the intramolecular interactions of the fuzzy region might compete with the intermolecular interactions of the binding element [51]. Physiologically, the co-existence of functionally different conformations may enable a global regulatory protein to be simultaneously engaged in multiple pathways. Alternatively, heterogeneous conformational ensembles may lead to specific context-dependent biomolecular recognition patterns via alternative interactions in response to environmental cues [52].

## Materials and Methods

### Strains

Strains used in this study are derivatives of S288C [53]. The *med15* deletion strain (OY320) is from the deletion collection [54] and was used throughout. A second *med15* knockout strain (JF1368) was used in the hGR reporter experiments (Fig. 1C). Phospho-site mutant strains were a gift from Dr. Patrick Cramer and Mathias Mann [38] and are also S288C derived. The genotypes of additional strains used in this study are listed in Table S1.

### UAS_G_-*lacZ* Strains

*GAL4 gal80* (JF2626) and *GAL4 GAL80* (JF2624) strains with an integrated UAS_G_-*lacZ* reporter were prepared by mating the *gal4 gal80* UAS_G_-*lacZ* strain MaV103 (MATa) [55] with BY4716 (MATα) [54]. Diploids from this cross were sporulated and tetrads were analyzed to identify *GAL4 GAL80* UAS_G_-*lacZ* and *GAL4 gal80* UAS_G_-*lacZ* strains. *GAL4* strains were identified by growth on YPGal media. Strains with *lacZ* reporters were identified by production of blue color on X-Gal plates. *GAL80* strains were identified by white colony color on X-Gal plates with glucose as the carbon source and blue colony color on X-Gal plates with galactose as the carbon source.

*MED15* was deleted from both *GAL4* UAS_G_-*lacZ* strains to produce *GAL80 med15* (JF2629) and *gal80 med15* (JF2631) strains. A *med15Δ*::kanMX4 deletion cassette was amplified from genomic DNA isolated from a *med15Δ* strain (OY320) using the primers *MED15* F-245 and *MED15* R+3498. The deletion cassette was transformed into JF2624 and JF2626 using a standard lithium acetate transformation protocol with the addition of a two-hour outgrowth in YPD. Transformation reactions were plated on YPD + 350 μg/mL G418. Deletion of the *MED15* locus was verified by PCR (primers *MED15* F-245 and *MED15* R+3498) of individual transformants.

### Variant Med15 Constructs

Plasmids constructed or acquired for this study are listed in Table S2. A series of constructs encoding synthetic Med15 proteins varying in number of domains and polyglutamine tract lengths were generated (Fig. 1). These constructs fall into four categories: internal deletions, KIX277-799Δ-Med15, KIXQ2Q3Δ-Med15, and KIXΔ-Med15, described below.

### *MED15* Internal Deletions

Two independent internal deletion mutants (153-687Δ and 46-619Δ) were prepared by restriction digestion of plasmid-borne intact *MED15* alleles. Designations correspond to amino acid residues removed. The 153-687Δ plasmid (pDC2285) was prepared by digestion with *Bsu*36I and subsequent yeast homologous recombination between sequence in the Q1 and Q3 tracts. The 46-619Δ internal deletion plasmid (pDC2286) was prepared by digestion with *Eco*RI and subsequent ligation to join compatible ends.

### *MED15* Gene Fragments (gBlocks)

gBlocks (Integrated DNA Technologies) corresponding to the KIX domain (bp 1-348), Q2 region (bp 841-1650), Q3 region (bp 1651-2394), and combined Q2 Q3 region (bp 841-2394) were synthesized. In gBlocks containing the Q2 region the Q2 tract and adjacent sequence (bp 1234-1446) were removed and replaced with an *Afe*I site by incorporating the silent base change, T1233C. In gBlocks containing the Q3 region the Q3 tract and adjacent sequence (bp 2002-2115) were removed and replaced with a *Bmg*BI site by incorporating the silent base changes T2001C and G2118C. All gBlocks have 80 bp of 5’ and 3’ homology to KIX277-799Δ-*MED15* on the pRS315 M-WT plasmid. Each gBlock was A-tailed using Taq DNA polymerase (New England Biolabs), ligated into pCR2.2-TOPO (TOPO, Invitrogen) and screened on LB-Amp + X-Gal plates. Light blue and white transformants were screened by PCR using M13 F and M13 R primers. For gBlock clones that included the Q2 and/or Q3 regions internal *MED15* primers (*MED15* F+1204 and *MED15* F+1924) were used in addition to the M13 primers to reduce the size of the amplified fragment. Finally, the plasmids were confirmed by sequencing and each clone was stored (GC1012, GC1013, GC1014, and GC1025).

A Q1 region (Q1R; bp 346-831) fragment was constructed with the Q1 tract replaced by an *Afe*I site using two step PCR. First, sequence on either side of the Q1 tract was amplified with overlapping forward and reverse primers having an *Afe*I site between Q1 flanking sequence (0Q F and 0Q R) paired with reverse (Q1R gap repair R) and forward (Q1R gap repair F) primers respectively. Next, the two PCR products and the outer forward and reverse primers were used to amplify a 0-Q1:*Afe*I Q1R fragment. This construct was ligated into the TOPO vector for storage and propagation (pDC2178).

gBlock DNA was digested out of the TOPO vectors using *Bst*XI (New England Biolabs), sites which are present in the vector on either side of the insert and are absent from gBlocks. The gBlock band was extracted and purified from agarose gels using the QIAquick Gel Extraction Kit (Qiagen). Alternatively, the KIX gBlock was PCR amplified from the TOPO vector using KIX gBlock F and KIX gBlock R primers.

### KIXQ2Q3Δ-Med15 Variants

KIXQ2Q3Δ-*MED15* constructs are intermediate in size relative to the KIX277-799Δ-*MED15* and full length *MED15* in that the KIX domain is still absent as it is in the KIX277-799Δ-Med15 while the central Q-rich region of Med15 is present (Fig. 1). These constructs were made by the addition of the Q2Q3 gBlock to either the 0-Q1 or 12-Q1 KIX277-799Δ-Med15 construct. The Q1R region of pRS315 M-WT, 0-Q1 KIX277-799Δ-*MED15*, and 12-Q1 KIX277-799Δ-*MED15* were PCR amplified using Q1R gap repair F and Q1R gap repair R. These PCR products were individually co-transformed with the Q2Q3 gBlock and *Pst*I digested pRS315 M-WT. The DNA fragment pairs have sequence overlap allowing for homologous recombination with each other as well as with the gapped plasmid. Two versions of 12-Q1 KIXQ2Q3Δ-*MED15* were constructed by using either pRS315 M-WT Q1R PCR (pDC2149) or using 12-Q1 KIX277-799Δ-*MED15* Q1R PCR (pDC2138). 0-Q1 KIXQ2Q3Δ-*MED15* (pDC2136) was constructed by using 0-Q1 KIX277-799Δ-*MED15* Q1R PCR.

Variant Q1 KIXQ2Q3Δ-*MED15* constructs were prepared by first introducing natural or synthetic Q1 sequences into the *Afe*I site in pDC2178 and then gap-repairing partially *Afe*I digested pDC2136 with a Q1R PCR product amplified from the pDC2178 derivative using the primers Q1R gap repair F and Q1R gap repair R. Natural Q1 tract length variants (19-Q1, pDC2150; and 21-Q1, pDC2151) were constructed by amplifying the Q1R sequence from pooled genomic DNA from multiple wine yeast strains using Q1R gap repair F and Q1R gap repair R primers. Synthetic Q1 tract length variants (12-Q1, pDC2149; 24-Q1, pDC2144; 36-Q1, pDC2146) were constructed by ligation of glutamine tract coding duplex DNA (5’-CAACAACAACAACAACAACAGCAGCAGCAGCAACAG). This method produced tracts of glutamine codons in multiples of 12. This method also produced tracts of polyleucine when the duplex was ligated in the reverse orientation (12L-Q1, pDC2293). A synthetic variant (47-Q1, pDC2147) was constructed by mismatch recombination between two tracts of 36 glutamine codons. A short Q tract (6-Q1, pDC2260), a non-Q spacer (Spacer-Q1, pDC2185), modified Q tracts (12PQ-Q1, pDC2262; 24PQ-Q1 pDC2263; 15LQ-Q1, pDC2291), and heterologous sequences (NAB3-Q1, pDC2292; FrHs-Q1, pDC2294; RvHs-Q1, pDC2295) were constructed by ligation of duplexed oligos. Duplexed oligos used in the ligation included: a glycine rich spacer sequence (5’-CCAGGTTCTGCTGGTTCTGCTGCTGGTGGT: PGSAGSAAGG); a short Q tract of 6Q (5’-CAACAACAACAACAGCAG); a coiled-coil disrupting sequence (5’-CCTCAACAACAGCCTCAGCAACCACAGCAACAACCA: PQQQPQQPQQQP); a coiled-coil promoting sequence (5’-TTGCAACAACAGTTACAGCAATTGCAGCAACAACTGTTGTTGCAA: LQQQLQQLQQQLLLQ); the C-terminal helix of Nab3 (5’-AATGTTCAAAGTCTATTAGATAGTTTAGCAAAACTACAAAAG: NVQSLLDSLAKLQK); and a portion of the human Med15 Q tract (5’-CTGCAGCTCCAGCAGGTGGCGCTGCAGCAGCAGCAGCAACAGCAGCAGTTCCAGCA GCAG) ligated in the forward (LQLQQVALQQQQQQQQFQQQ) and reverse orientation (LLLELLLLLLLLQRHLLELQ).

### KIXΔ-Med15

Plasmid KIX277-799Δ-*MED15* (pRS315 M-WT) was digested with *Bst*API to create a gap between the Q1 containing region (Q1R) and the MAD region. The gap was repaired using the Q2/Q3 gBlock isolated from GC1014 to generate pDC2149 (KIXQ2Q3Δ-Med15). pDC2149 was sequentially digested at the Q3 locus using *Bmg*BI and gap repaired using the Q3 PCR fragment (primers *MED15* F+1682 and *MED15* R+2385) from the lab strain BY4742 and then digested at the Q2 locus using *AfeI* and gap repaired using the Q2 PCR fragment (primers *MED15* F+854 and *MED15* R+1781) from the lab strain BY4742 to generate pDC2217 (KIXΔ-Med15).

### Split Ubiquitin

#### CUb fusions

A P*_MET17_* C-ubiquitin R*URA3* plasmid (GC408; Addgene 131163, [56]) was linearized with *SalI* directly upstream of the C-ubiquitin sequence and gap-repaired with PCR products amplified from the OY235 yeast genome. To create MSN2_TAD_-CUb (pDC2279) the first 813 bp of the *MSN2* ORF were amplified using Msn2_TAD_ F+R. For KIX-CUb (pSL2311 and pSL2318) the insert was amplified using KIX-CUb F+R primers. Recombinant plasmids were confirmed by colony PCR.

NUb fusions: A P*_CUP1_* N-ubiquitin plasmid was constructed by amplifying the *CUP1* promoter through N-ubiquitin sequence from the integrating plasmid (Addgene 131169, [56]) and gap repairing it into pRS315 linearized with *Bam*HI. Subsequently the 3’UTR sequence from *MED15* was gap repaired into the plasmid linearized with *Hin*DII*I* downstream of the ORF to create pDC2278. Fragments of the *MED15* gene were amplified and gap repaired into pDC2278 linearized with *PstI* downstream of the N-ubiquitin sequence (KIX, pDC2283; Q1R, pDC2284; KIX+Q1R, pDC2282; Q1R-NAB, pDC2280; Q1R-Hs-rev, pDC2281; Q1R-0Q, pDC2301; Q1R-24Q, pDC2302; Q1R-Spacer, pDC2303; Q1R-6Q, pDC2304; Q1R-12PQ, pDC2305; Q1R-24PQ, pDC2306; Q1R-12L, pDC2307; Q2Q3R, pSL2310.

### Bacterial Expression Plasmids

Bacterial expression plasmids pET21a-mCherry-Med15 Q1R, pSL2272 and pET21a-mCherry-Med15 Q1R-0 (pSL2289) were constructed by Gibson Assembly. Desired inserts were amplified with Q5 DNA polymerase using the yeast genome or preexisting plasmids as templates (See primer list, Table S4). PCR products were purified using PCR clean-up kits (Qiagen). The inserts and backbones were assembled by NEBuilder HiFi DNA Assembly Cloning Kit in a 10ul reaction. 2 µL was transformed into *E.coli* DH5α. Plasmids were isolated and confirmed by DNA sequencing.

### Yeast Methods

#### Colony PCR

Yeast colonies were used for rapid PCR screening using Taq polymerase, colony PCR buffer (final concentrations: 12.5 mM Tris-Cl (pH 8.5), 56 mM KCl, 1.5 mM MgCl_2_, 0.2 mM dNTPs), and primers at a final concentration of 0.2 μM. Reaction mixes were aliquoted into PCR tubes and then a small amount of yeast cells were transferred from a streak plate into each tube using the end of a 200 μL micropipet tip. A standard hot-start thermocycler program with extension at 68°C was adjusted for the Tm of the primer set and size of the amplification product.

#### Yeast Transformation

Transformations of *med15Δ* strains were conducted using the Frozen-EZ Yeast Transformation II kit (Zymo Research) with modifications. 10 mL of early log phase cells were pelleted and washed with 2 mL EZ 1 solution, repelleted and resuspended in 1 mL EZ 2 solution. Aliquots were frozen at -80°C. 0.2-1 μg DNA and 100 μL EZ 3 solution were added to 10 μL competent cells for each transformation. Transformations were plated on selective media following incubation at 30°C for 45 minutes. Transformations into all other strains were conducted using a standard lithium acetate transformation protocol [57, 58].

#### Media and Phenotype Testing

Cultures were grown in rich media (YPD), synthetic complete media lacking single amino acids (SC-Leu or SC-Ura), or synthetic defined media supplemented with required amino acids (SD+Lys). Synthetic media types described using one- or three-letter amino acid abbreviations. Spot assays were performed on rich media or synthetic complete media with additives.

Exponentially growing subcultures were diluted differentially to achieve a consistent initial concentration of 5x10^6^ cells/mL. 5-fold or 10-fold serial dilutions were carried out in the wells of a sterile 96-well microtiter dish. 2 μL volumes of each dilution were spotted on different types of solid media. Plates were incubated at desired temperatures (22°C, 30°C, and 38°C). Phenotypes were observed and imaged daily using a flatbed scanner. Images were selected for figures based on the incubation times at which differential phenotypes on specific media types were most apparent compared to growth on YPD at the same time. In some cases (various concentrations of ketoconazole, even the wild type strain was affected.

The results of spot assays were visualized using ImageJ [59] (Fig. 4). Grayscale TIFF formatted images of plates were imported into ImageJ. Background pixels, that did not correspond to yeast colonies, were set to an intensity of 0 using an intensity threshold cutoff. Threshold cutoffs determined for each plate separately. An area containing yeast colonies was selected using the rectangle tool. The same rectangular area selection was used across a single experiment. For each strain the percent area was measured (pixels corresponding to colonies/total area * 100) and normalized to the growth of that strain on YPD.

A microtiter dish based chemical sensitivity assay was implemented as previously described [60] with minor modifications to achieve increased resolution. Cells of each genotype in biological triplicate (transformants) were grown to saturation in SC-Leu media, diluted to an OD_600_ of 0.1-0.2 and inoculated 1:1 into 2xYPD with various concentrations of drug or in alternative types of media and allowed to incubate overnight at the indicated temperature. OD_600_ readings were conducted using a microplate reader following 30 seconds of shaking at an endpoint of 20-24 hours. Relative growth was calculated after background subtraction by dividing the absorbance for each treated well with the corresponding value of the 0-drug control well. Relative growth for each genotype was plotted for a single drug concentration chosen for causing a reduction in growth of the wild type *MED15* strain of less than 50%.

### β-Galactosidase Reporter Assays

#### Human Glucocorticoid Receptor Tau 1 Fragment Dependent Reporter

Yeast carrying an expression vector for the human glucocorticoid receptor Tau 1 fragment transcription factor (hGRτ1) fused to LexA and the LexA-driven *lacZ* reporter (JF2768) were transformed with a third plasmid carrying the desired *MED15* allele or an empty vector (pRS315). As previously described, transformants were prescreened on X-Gal plates to identify transformants displaying the “average” amount of reporter activity for that strain [18]. Outliers that were either noticeably more or less blue than the other transformants were excluded.

At least 3 transformants per strain were grown in minimal media (SD+AdeLys) to a concentration of approximately 2x10^7^ cells/mL. Induction of the hGR transcription factor was by addition of CuSO_4_ to 0.25 mM for 1 hour. Cultures were pelleted and stored at -80°C in the presence of glass beads and protein extraction buffer. Protein extracts were prepared and analysis conducted within a week to avoid degradation or deterioration of the sample.

#### Gal4 Dependent Reporter

*med15Δ gal80Δ* strain (JF2631) containing an integrated UAS_G_-*lacZ* reporter was transformed with plasmids containing the desired *MED15* allele or an empty vector. Transformants were subcultured in SC-Leu 2% raffinose and grown at 30°C to log phase (1-2x10^7^ cells/mL). *med15Δ GAL80* strains (JF2629) containing an integrated UAS_G_-*lacZ* reporter were likewise cultured in SC-Leu 2% raffinose to saturation but then subcultured in SC-Leu with 2% raffinose + 2% glucose or SC-Leu 2% raffinose + 2% galactose and grown at 30°C to log phase (1-2x10^7^ cells/mL). Cultures prepared for protein extraction as above and stored at -80°C.

#### Protein Extraction

Yeast cells were harvested from log-phase cultures in reporter-specific growth conditions and frozen at -80°C. Pelleted yeast cells were resuspended in Breaking Buffer (0.1 M Tris pH 8, 20% glycerol, 1 mM DTT) in tubes containing glass beads (200 mg, 0.4 mm) and were lysed using a TissueLyser LT (Qiagen). To minimize protein degradation, 5 μL of 40 mM PMSF was added periodically throughout the process of preparing extracts. Tubes were alternately shaken at 50 oscillations/second for 20-30 seconds and then placed on ice for at least 1 minute for a total of 2 minutes of shaking. Cell debris was pelleted at 13,000 rpm for 10 minutes at 4°C. The supernatant was transferred to a new tube containing additional PMSF. Protein extracts were stored at -80°C or kept on ice until used for subsequent analyses.

### Protein Concentration Determination (Bradford Assay)

Total protein concentrations were determined using a modified Bradford Assay (BioRad). 2-10 μL of each protein extract or BSA standard (100-1000 mg/mL) was added in duplicate wells of a 96-well plate. 200 μL of Bradford assay working solution was added using an 8-tip multichannel pipet. A standard curve was generated using a series of BSA standards of known concentrations designed to produce absorbance readings within the linear range of the spectrophotometer. Final absorbance readings were taken after 15 minutes at a wavelength of 595 nm. The average absorbance of blank wells was subtracted from all readings. A best fit linear regression based on the standard calibration curve was used to determine sample concentration.

### Enzymatic Assay

Approximately 10 ng of total protein (5-50 μL protein extract) was diluted to 1 mL in Z-buffer (60 mM Na_2_HPO_4_, 40 mM NaH_2_PO_4_, 10 mM KCl, 1 mM MgSO_4_, 50 mM β-mercaptoethanol (BME)) in 13 mm glass tubes and incubated in a temperature block held at 28°C. 200 μL ONPG (4 mg/mL in Z-buffer; made fresh for each experiment) was added to each tube (time 0). Reactions were stopped once the solutions turned yellow by addition of 500 μL of 1 M Na_2_CO_3_. A_420_ measurements were taken for each sample.

### Data Processing

Beta-galactosidase values were calculated using the formula:

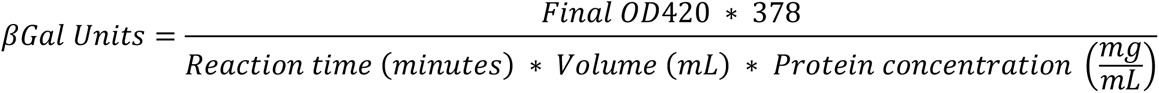

Due to systematic difference between experiments, such as different ONPG solution preparations or quality of extracts, relative expression values were calculated for each experiment to be able to compare strains measured in different experiments. When necessary, the average β-gal value for the wild type strain was set to 100% for each experiment.

### Split Ubiquitin Assay

Yeast strains containing both an N-ubiquitin and a C-ubiquitin-R*URA3* expressing plasmids were cultured in SD+LysUra or SC-HisLeu to log-phase at 30°C to a concentration of ∼1x10^7^ cells/mL. CuSO_4_ was added to a final concentration of 0.05-0.25 mM and the cultures incubated for an additional hour. In some experiments, methionine was added to 20 µM during this incubation. The addition of small amounts of methionine clarified the phenotype on plates lacking uracil. Cultures were serially diluted and 2 µL was plated on media lacking methionine and uracil as well as media containing 5-FOA [61].

### RNA Methods

Yeast strains were cultured in SC-Leu to saturation and then subcultured in 10 mL YPD and grown to log-phase at 30°C to a concentration of ∼2x10^7^ cells/mL. After incubation yeast were pelleted, washed with cold water, frozen with dry ice, and stored at -80°C. RNA was extracted using a hot acid phenol protocol with modifications [62]. Pellets were resuspended in 400 µL TES solution (10 mM Tris, 10 mM EDTA, 0.5% SDS) and an equal volume of acid phenol was added to each tube and vortexed for 10 seconds. Tubes were incubated in a 65°C water bath for 1 hour and vortexed every 10-12 minutes. Phases were separated by microcentrifugation in the cold for 5 minutes. The aqueous layer was extracted into a new tube avoiding the DNA-enriched interface. A second round of acid phenol extraction was conducted followed by a chloroform extraction. RNA was precipitated with 300 µL of 4 M LiCl in dry ice for 20 minutes followed by 5 minutes of high-speed centrifugation at 4°C. The pellet was washed with 500 µL ice cold 70% ethanol and then dried for 10 minutes at 37°C. RNA pellets were resuspended in DEPC treated water and stored at -20°C.

Contaminating genomic DNA was removed using the DNase Max Kit (Qiagen). 10 µL of a 1x Master Mix (5 µL 10X Buffer, 5 µL water, and 0.5 µL DNase I) was added to 40 µL RNA. Tubes were incubated at 37°C for 30 minutes. DNase was removed by addition of 5 µL of the DNase removal resin and room temperature incubation for 10 minutes with periodic agitation followed by centrifugation. Alternatively, DNase was removed using an RNA clean-up column (Zymo Research). The concentration of the purified RNA was determined using a NanoDrop spectrophotometer (Thermo Scientific).

### cDNA First Strand Synthesis

cDNA was prepared using Superscript III First-Strand Synthesis System (Invitrogen) or Lunascript (New England Biolabs). Transcripts were amplified with either random hexamer (Invitrogen) or anchored oligo-dT20 (Integrated DNA Technologies) primers. For anchored oligo-dT20 primers 1 μg RNA, 50 ng primer, and 0.01 μmol dNTPs were mixed in a final volume of 10 μL. Tubes were incubated for 5 minutes at 65°C and then 2 minutes on ice. 10 μL cDNA Master Mix (2x RT buffer, 10 mM MgCl_2_, 0.02 M DTT, 40 U RNase OUT, 200 U Superscript III Reverse Transcriptase) or mock Master Mix which did not contain reverse transcriptase was added to each tube which were then incubated for 60 minutes at 50°C and 5 minutes at 85°C. To degrade RNA, 2 U RNase H was added to each tube and then the tubes were incubated for 20 minutes at 37°C. For random hexamer primers tubes were incubated 10 minutes at 25°C and then 50 minutes at 50°C.

### qRT-PCR

Transcript abundance was quantified for each sample relative to a normalization transcript using the PerfeCTa SYBR Green FastMix (Quantabio) in 96-well PCR plates (Hard-Shell 480 PCR Plates, Bio-Rad) measured in a LightCycler 480 (Roche). Individual 10 µL reactions consisted of 5 µL FastMix, 1 µL target-specific primer pairs (0.5 µL 10 µM forward primer and 0.5 µL 10 µM reverse primer), 3 µL DEPC treated water, and 1 µL cDNA, mock cDNA, or water. Plates were sealed with optically clear film (PlateSeal). A standard SYBR green PCR program was used with modifications: 95°C for 5 minutes, 45 cycles of 95°C for 10 seconds, 55°C for 10 seconds, and 72°C for 20 seconds with a single fluorescence acquisition. A melting curve was conducted directly following the PCR program to confirm the presence of individual species amplified in each well. The melting curve program was 95°C for 5 seconds, 65°C for 1 minute, and ramp up to 97°C by 0.11°C/s with continuous fluorescence acquisition. *ALG9*, a gene encoding an alpha 1,2 mannosyltransferase, which is known to be stably expressed across growth conditions in yeast [63], was used as a normalization transcript. All samples were measured with three technical replicates. Mock cDNA (prepared without the addition of reverse transcriptase) and water were used as negative controls.

### Data Analysis

The relative abundance of target transcripts was determined by calculating the crossing point (CP) or cycle threshold (CT) value for each reaction. CP values were calculated using the Second Derivative Maximum method implemented in the LightCycler 480 Software (Roche). CT values were calculated using the Comparative CT (ΔΔCT) method implemented in the QuantStudio 3 Software (Applied Biosystems). The average CP or CT for technical replicates amplifying the target transcript with a specific RNA sample was normalized to the average CP or CT for technical replicates amplifying the normalization transcript. This ratio was then compared across samples as a depiction of relative abundance of the target transcript in each sample.

### Primers

Primers used in this study are listed in Table S3. All primers were synthesized by Integrated DNA Technologies. The use of each primer in construction, screening, and sequencing is noted in Table S3a, b and c. All primer pairs used to amplify transcripts in qRT-PCR are listed in Table S4. Primer pair efficiency was measured by qRT-PCR analysis of serial dilution of control RNA.

### Protein Expression and Purification

All bacterial expression plasmids were transformed into *E. coli* BL21-AI. Transformants were incubated in 250mL LB media with 0.1% glucose and grown to log phase at 37°C. 0.5mM IPTG and 0.2% arabinose were added to the media and the bacteria cultured at room temperature overnight. Bacteria were harvested by low-speed centrifugation for 20 minutes. Pellets were resuspended in 10 mL buffer A (50 mM pH7.5 Tris-HCl, 500 mM NaCl) with PMSF added to 1 mM. Bacterial lysis was by sonication. The lysate was cleared by centrifugation at 3000 rpm for 30 minutes at 4°C

Gcn4 and Med15 supernatants were mixed with 5 ml Ni-NTA beads (Qiagen) and incubated at 4°C for 1 hour. The beads were spun down at 600 rpm and the supernatant removed. Beads were washed 2x with 5 ml buffer A containing 50 mM imidazole, and then eluted with 5 ml buffer A containing 250 mM imidazole.

For Msn2, a 1 ml HisPur™ Cobalt (Thermo Fisher) column was equilibrated with 2 ml of buffer A prior to the addition of the bacterial lysate and a 1x wash step with 4 ml buffer A. Elution was with 1 ml buffer A + 250 mM imidazole. The eluate was diluted with 9 ml buffer W (100 mM Tris-HCl pH 8.0, 150 mM NaCl, 1 mM EDTA) and added to a 1 ml Strep-TactinXT resin (IBA Lifesciences) column pre-equilibrated with 2 ml buffer W. An initial wash with 5 ml of buffer W was followed by elution with 1 ml buffer W + 50 mM biotin.

For Med15 Q1R with a 12L Q1 insert, 2% final concentration of SDS was added to the lysis buffer to solubilize the protein. After sonication and centrifugation, KCl was added to the supernatant at the final concentration of 400 mM. SDS-KCl crystal were allowed to form for 1 hour at 4°C after which these were removed by centrifugation at 3000 rpm for 30 minutes. The supernatant was purified by Ni-NTA chromatography as described above. After elution, KCl was added to the elution at the final concentration of 400 mM and incubated for 1 hour at 4°C again followed by centrifugation at 13000 rpm for 5 minutes to ensure maximum SDS removal. The supernatant was kept for further usage.

All purified proteins were dialyzed 3 times, 1 hour each, in 700 ml buffer K (50 mM Tris-HCl pH 7.5, 200 mM KCl, 20 mM NaCl, 30 mM MgCl_2_, 1 mM DTT, 10% Glycerol), and concentrated using a Millipore 30,000 MWCO centrifuge concentrator to reduce the volume to 100 ul or less. The concentration of the protein was measured spectrophotometrically using a NanoDrop One (Thermo Scientific) and purity confirmed by SDS-PAGE analysis followed by Coomassie Blue staining.

### *In vitro* droplet formation and turbidity assays

Purified mCherry-Med15 and Gcn4-GFP were mixed to a final concentration of 20 μM (10 μM each protein) in 5 µL. 5 µL of crowding buffer (6% PEG-8000 final concentration) [30] was added and mixed thoroughly. 7µL was pipetted onto a cover slip and allowed to stand for at least 1 minute before being covered with a microscope slide. Slides were observed using a Zeiss fluorescence microscope. 1,6 hexanediol was added at the start of reactions at the indicated concentration.

## Acknowledgements.

The authors gratefully acknowledge the generous gift of strains and plasmids from Dr. Patrick Cramer and Mathias Mann, (phospho-site mutant strains), Dr. Young Chul Lee (glucocorticoid reporter plasmids), and Nils Johnsson via Addgene (split ubiquitin vectors) and Dr. Richard Young for the GFP-Gcn4 expression vector, RY8740. The authors also appreciate comments and suggestions by Dr. Daniel Weeks and Dr. Bryan Phillips, as well as members of the lab, and anonymous reviewers.

